# Conserved allomorphs of MR1 drive specificity of MR1-restricted TCRs

**DOI:** 10.1101/2023.07.17.548997

**Authors:** Terri V Cornforth, Nathifa Moyo, Suzanne Cole, Emily Lam, Tatiana Lobry, Ron Wolchinsky, Angharad Lloyd, Katarzyna Ward, Eleanor M Denham, Gurdyal S Besra, Natacha Veerapen, Patricia T Illing, Julian P Vivian, Jeremy M Raynes, Jérôme Le Nours, Anthony W Purcell, Samit Kundu, Jonathan D Silk, Luke Williams, Sophie Papa, Jamie Rossjohn, Duncan Howie, Joseph Dukes

## Abstract

Major histocompatibility complex class-1-related protein (MR1), unlike human leukocyte antigen (HLA) class-1, has until recently been reported to be monomorphic. Tumor cell-specific MR1 restricted T cell receptors (TCRs) have been described, offering potential therapeutic application for cancer treatment. We show that human T cells expressing a TCR derived from an MR1-restricted T cell clone, termed MC.7.G5 (7G5.TCRT), retain MR1-directed cytotoxicity. However, activity is not pan-cancer, as initially reported with the clone MC.7.G5. Recognition is restricted by an allelic variant of MR1 (MR1*04) which is present at approximately 1% of the population at the heterozygote level. The 7G5 TCR is not cancer specific, as 7G5.TCRT and 7G5.TCRT-like TCRs react to both cancer and healthy cells expressing MR1*04 alleles. These data demonstrate that healthy individuals can harbor T cells reactive to an MR1 variant displaying self-ligands expressed in cancer and benign tissues. Targeting MR1 in cancer will require identification of cancer-specific presented ligands, and careful confirmation of cancer specificity of TCRs. MR1*04 may behave as an alloantigen warranting further study.

## Introduction

The MHC class I-related protein 1 (MR1) is one of several non-classical human leukocyte antigen (HLA) molecules ^1–8^ Functionally, MR1 presents antigens from folate and riboflavin-derived metabolites to the immune system, some of which are derived from bacteria or yeast. Unlike the classical polymorphic HLA Class I molecules, which present peptide antigens (pHLA), MR1 was widely considered to be monomorphic. However, in 2021 Rozemuller et al identified five distinct MR1 allele group variants in a series of 56 DNA samples taken from cells with diversity in HLA^9^. They adopted MR1*01 as nomenclature of the wild type allele with MR1*02, the most common variant at 21% in their analysis, demonstrating a single nucleotide polymorphism (SNP) at H17R in the parent protein. MR1*04 has both an R9H and H17R polymorphism and has a heterozygous frequency of approximately 1 in 100 Caucasians.

Structural studies of MR1 molecules demonstrate a similar overall architecture to HLA class I molecules, formed of a heavy chain composed of α1 and α2 helices forming the binding groove and an α3 domain non-covalently bound to β2m^10, 11^. The binding groove of MR1 has an A’ and F’ pocket. The A’ pocket binds ligands and is lined with aromatic and basic residues. These residues create an environment permissive for binding of microbially derived vitamin B12 metabolites and restrict the space available for binding to peptides^2, 10, 11^ . The two basic residues in the A’ pocket, R9 and K43, are critical for interaction with ligands. R9 is conserved across mammalian species and interacts directly with ligands^12^. K43 is essential for formation of covalent Schiff bonds with many pyrimidine based ligands^1, 12–14^

Many aspects of MR1 biology have been revealed by studies of a unique class of T cells, known as mucosal-associated invariant T (MAIT) cells, which recognize ligands derived from bacterial metabolites bound to MR1 using a limited (invariant) set of αβ and γδ T cell receptors (TCRs)^3, 15–20^. The nature of the ligands presented by MR1 to MAIT cells has been the subject of numerous investigations^1, 21–23^. Interestingly, the antigens and TCRs are conserved sufficiently such that there is cross-reactivity between human and murine derived MR1s and MAIT cells. The invariant nature of MAIT TCRs indicates that this class of T cell operates at the interface between adaptive and innate immunity.

A distinct subset of MR1-restricted T cells (MR1T) have been identified with the apparent ability to specifically recognize and kill cancer cells^24–26^ . The MR1 ligands recognized by cancer specific MR1T remain poorly understood but appear distinct from those of MAIT cells^26, 27^ . Within the MR1T cell family there are patterns of ligand preference illustrated by a differential dependence for recognition on the K43 residue in the MR1 binding groove^25^. Isolation and study of TCRs from putative MR1T present an attractive potential source of novel cancer therapies. Targeting peptides presented by specific HLA Class I molecules (e.g., HLA-A*02:01, HLA-B*08:01, HLA-C*07:01, etc.) somewhat limits broad utility of a TCR based therapy as they must be custom-developed for use in limited fractions of the human population. The reported monomorphic nature of MR1 raised the possibility that therapies directed towards MR1-displayed antigens have potential to be effective across the entire human population. Yet, recent reverse translation from a patient with a pervasive and unusual chronic infective phenotype, resulted in the identification of a single nucleotide polymorphism in the patient’s MR1 for which they were homozygous^12^ . The point mutation was identified at R9H of the mature MR1 protein, within the antigen binding groove of MR1.^26, 27^ The investigators observed that MAIT cells were completely absent in the patient’s circulating peripheral blood mononuclear cells (PBMC), demonstrating functional outcomes of polymorphism of MR1.

We undertook to express TCRs isolated from MR1T clones in polyclonal human T cells to test the hypothesis that these TCRs could have pan-cancer therapeutic utility. Validation of our approach led to the understanding that conserved SNPs present in MR1 can drive T cell activation, in a highly MR1*04-specific manner, that is not exclusive to tumor cells^9, 12^. Collectively these data highlight the need for deeper understanding of MR1 biology in the context of cancer and raise the possibility that MR1 polymorphism may need to be considered in the context of allotransplantation and graft versus host disease (GvHD).

## Results

### The TCR from MC.7.G5 is functionally comparable in TCRT format to clinically validated pHLA directed TCRTs

A T cell clone, MC.7.G5, has been described to be MR1-restricted, have pan-cancer reactivity, not recognize normal cells, or cells subject to various forms of cellular stress^25^. To test the potential of the MC.7.G5 TCR for cancer therapeutic translation the MC.7.G5 TCR was cloned into a lentiviral vector for transduction into human T cells (Figure 1a, Supplemental Figure 1). Transduction of the native TCR sequence derived from MC.7.G5 (“nat.7G5”) resulted in poor cell surface expression in primary human T cells (expressing endogenous TCRs) (data not shown). Previous studies have shown that murinization of TCR constant domains and substitution of hydrophobic residues near the transmembrane domain enhance cell surface expression whilst maintaining TCR specificity^28^. To test if this strategy would enhance cell surface expression of the MC.7.G5-derived TCR on T cells we murinized the constant domains of the vectored TCR sequence and introduced hydrophobic residues into the transmembrane portion of the alpha chain. Locations of the hydrophobic residues are depicted in Supplementary 1a. This resulted in enhanced expression of the murinised MC.7.G5- derived TCR (“7G5”) at the cell surface (Supplementary Fig 1b) and was consistently detected in 20-50% of healthy donor CD8+ T cells (Figure 1b). Transduced T cells, henceforth called ‘7G5.TCRT’ cells, secrete IFNγ in an MR1-restricted fashion in response to the MR1-positive non-small cell lung cancer line A549, and not to CRISPR MR1 knock out (ko) A549 cells (Figure 1c). When compared to TCRs in clinical development (c259 and CD8a.c259 TCRs recognizing NY-ESO-1^29^, ADP-A2M4 and CD8a.ADP-A2M4 TCRs recognizing MAGE-A4^30^ and IMA203, recognizing PRAME^31^, 7G5.TCRT cells demonstrate comparable activity as demonstrated by IFNγ production, and cytotoxicity (Figure 1d). These results confirm that MR1-restricted 7G5.TCRT can exhibit comparable potency to affinity-enhanced TCRT which are restricted to HLA class I targets and have been demonstrated to have anti-tumor activity in the clinic.

**Figure 1.**
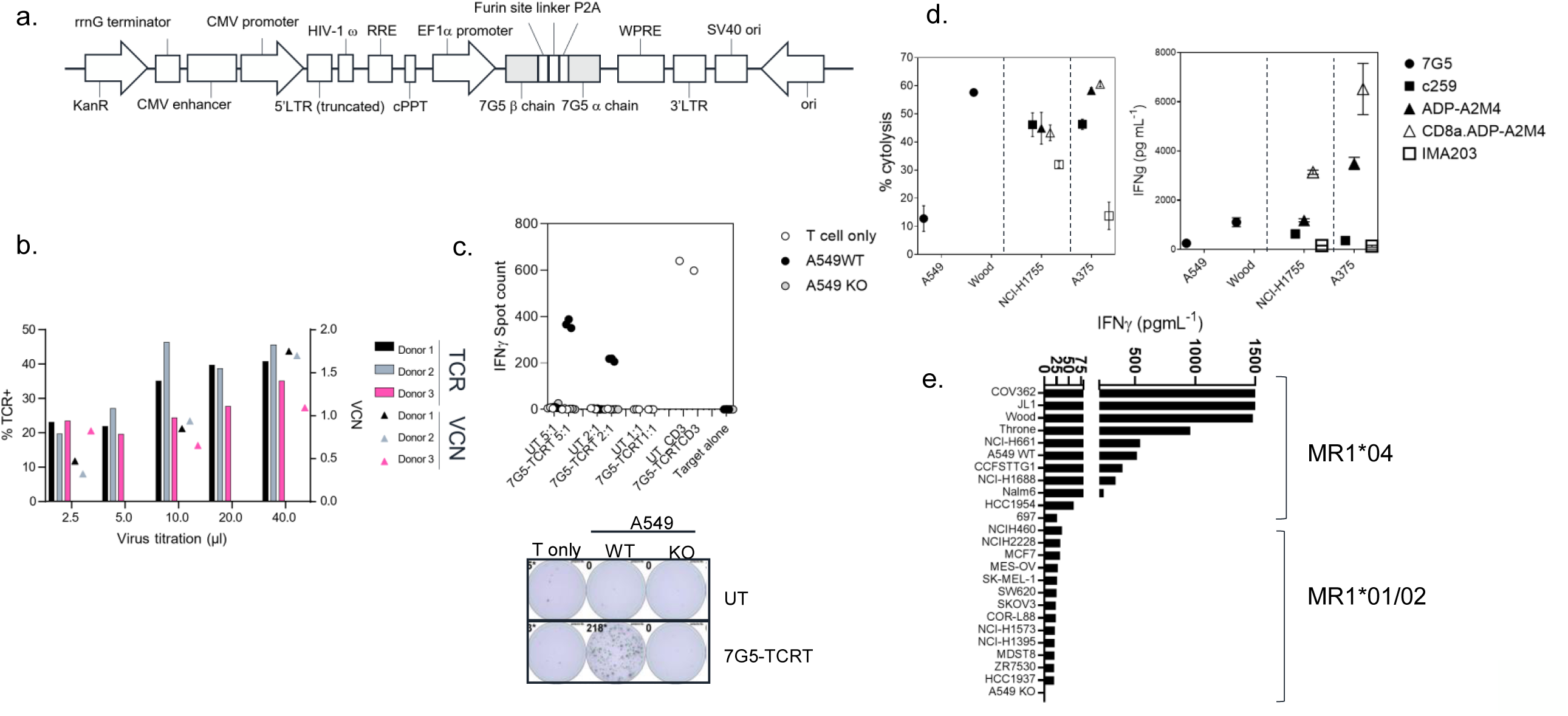
7G5-TCRT preferentially targets cancer cells bearing allelic variant MR1*04. **1a** Features of the lentiviral vector (LVV) used to transduce the 7G5 TCR into primary T cells. The 7G5 TCR used murine constant regions to enhance chain pairing and cell surface expression in primary T cells. A detailed map is shown in Supplementary Figure 1. **1b** Transduction efficiency of 7G5-encoding LVV in healthy CD8 T cells derived from PBMC. Staining of 7G5-TCRT for TCRBV25 was used to assess 7G5 expression. X axis indicates volume of LVV supernatant used per transduction. Right Y axis indicates viral copy number (VCN). **1c** Potency of 7G5-TCRT against MR1-expressing NSCLC line A549 or A549 MR1 knockout cells. ELISpot assays were used to measure IFNγ secretion by 7G5-TCRT cells in response to the targets at the indicated effector to target ratios. Bottom panel: Representative IFNγ ELISpot showing reactivity of 7G5-TCRT to MR1 wild type or MR1 knockout A549 cells. The data shown in Figure 1c is representative of at least 6 biological replicates. **1d.** Comparison of 7G5-TCRT cytotoxicity and IFNγ release in response to cancer cell lines with the response of clinically relevant HLA-restricted TCRs to antigen-positive cancer cell lines. Left hand panel. Mean (±SD, n=3) percentage cytolysis above background of cancer cell lines (X axis) by human T cells transduced with the indicated T cell receptor after 48 co-culture (symbols). Cell numbers varied depending on target but Effector target ratios (E:T) remained at 5:1 Data is representative of two biological replicates. Right hand panel. Mean (±SD, n=2-9) IFNγ concentration measured by ELISA in the supernatant 18 hours following co-culture of 20,000 cancer cells l(X axis) and 100,000 human T cells transduced with the indicated T cell receptor (symbols). E:T ratio 5:1. Data is representative of two biological replicates. **1e** Reactivity of 7G5.TCRT to 25 cancer cell lines (of 135 analyzed, Supplementary Table 1) ranked according to reactivity as measured by IFNγ ELISA at 48 hours following co-culture of 20,000 target cells and 60,000 7G5.TCRT, E:T ratio 3:1

### 7G5.TCRT do not demonstrate pan-cancer reactivity

We next tested the hypothesis that the 7G5 TCR has pan-cancer recognition ability, in line with the reported activity of the parent CD8^+^ clone MC.7.G5^25^, by measuring reactivity of 7G5.TCRT to an extensive panel of cancer cell lines. We set up reactivity assays of 7G5.TCRT to 135 cancer cell lines, using IFNγ as a readout of activity. We observed convincing 7G5.TCRT reactivity, defined as >50pg/ml IFNγ, in co-culture with only 7% (9/125) of cancer cell lines (Figure 1e and Supplementary Table 1). We tested several hypotheses to establish the attributes required for cancer cells to be recognized by 7G5. Reactivity of 7G5.TCRT did not correlate with surface levels of MR1 (Supplementary Figure 2), number or polarization of mitochondria, levels of superoxide within the cell, cell cycle time, or transcriptional levels of the alternate MR1 transcripts A and B (data not shown). Based on a recent report of multiple polymorphic forms of MR1 we investigated whether specific allomorphs of MR1 were recognized preferentially by 7G5.TCRT^9^.

### The 7G5 T cell receptor preferentially targets cancer cells bearing the MR1*04 allele

To investigate whether 7G5.TCRT is preferentially restricted by previously described MR1 variants^9, 12^, we PCR-amplified and sequenced the MR1 gene from the 135 lines tested (Supplementary Table 1). In target cell lines utilized for comparison with 7G5.TCRT against clinical stage TCRs, A549, the breast carcinoma line “Wood” and lung adenocarcinoma line “Throne” are all heterozygous for the MR1*04 allele. 7G5.TCRT reacted comparatively poorly to lines A375 and NCI-H1755 which express MR1*01 but not MR1*04 (Figure 1c). Using IFNγ as a measure of reactivity of 7G5.TCRT to these cancer lines we showed definitively that 7G5.TCRT preferentially targets cells expressing the MR1*04 allele, while responses to over 100 MR1*01 and MR1*02 expressing cells were typically much weaker or non-existent. IFNγ production by 7G5.TCRT to MR1*04-bearing targets was between 2 and 110-fold greater than to those not expressing this allele (Figure 2d and supplementary table 1). The lack of activity of 7G5.TCRT against MR1*02 expressing cells shows that the H17R mutation is not sufficient for recognition of MR1*04.

**Figure 2.**
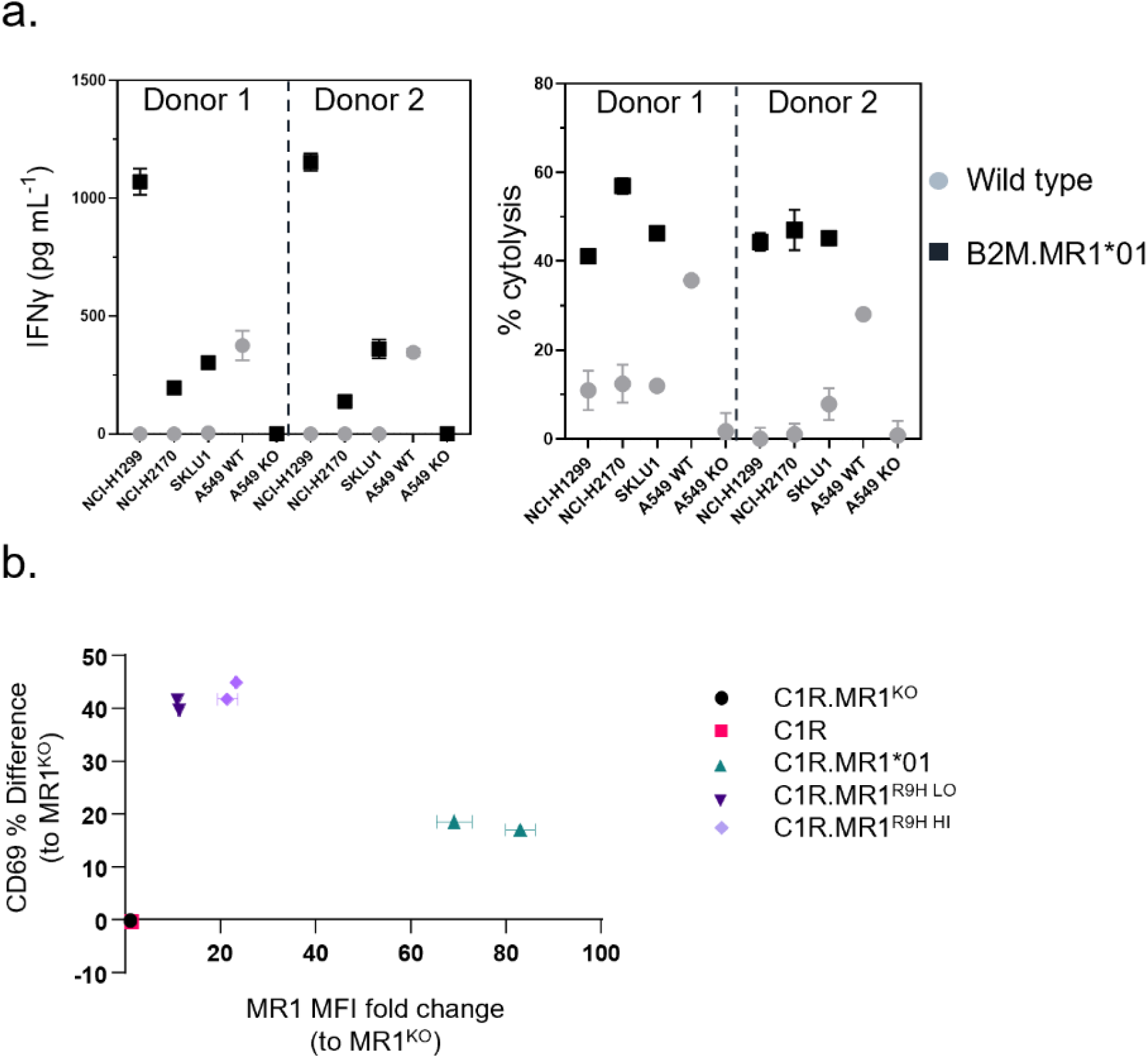
Over-expression of MR1*01 is required for reactivity of 7G5-TCRT to cells lacking MR1*04. **2a** Right panel panel: Mean (±SD, n=3) percentage cytolysis above background of WT or β2m.MR1- expressing cancer cell lines (X axis) by 7G5.TCRT cells from 2 donors after 24 hours co-culture. Cell numbers varied depending on target but Effector target ratios (E:T) remained at 5:1. Left panel: Mean (±SD, n=3) IFNγ concentration measured by ELISA in the supernatant 18-24 hours following co-culture of 20,000 WT or β2m.MR1-expressing cancer lines (X axis) and 100,000 7G5.TCRT, E:T ratio 5:1. MR1 haplotypes of the cells investigated; NCI-H1299 and NCI-H2170: MR1*01, SKLU1: MR1*02, A549: MR1*04/*01 heterozygous. **2b** Activation of Jurkat.β2m^ko^.7G5 measured as %CD69^hi^ (after subtraction of the mean %CD69^hi^ on stimulation with C1R.MR1^ko^ cells) vs fold change of the Median Fluorescence intensity of MR1 expressed by the stimulating APC compared to C1R.MR1^ko^. Data are from two independent experiments. Each point represents the mean (+/- SD) of three in-experiment replicates for CD69 and two in-experiment replicates for MR1.

### 7G5.TCRT only respond to MR1*01 on target cells at supra-physiologic expression levels

We next explored the possibility that preferential reactivity of 7G5.TCRT to MR1*04 allele expressing cancer cells might be due to MR1*04 being more abundantly expressed at the cell surface than MR1*01. We did not observe any correlation between surface expression of MR1 allomorphs on cancer cell lines, as measured by flow cytometry, and 7G5.TCRT efficacy, as measured by IFNγ release (Supplementary Figure 2). MR1*04 expression stimulates 7G5.TCRT to a much greater extent than MR1*01 when the levels of expression on the cell surface are comparable. Given most studies to date with MR1-reactive T cells rely on MR1*01 over-expressing lines as targets, we next investigated whether 7G5.TCRT could react to MR1*04 negative cancer lines engineered to express supra-physiological levels of the MR1*01 allele. The cancer cell lines NCI-H1299, NCI-H2170 and SKLU1, all negative for MR1*04 expression, were chosen as they are poor targets for 7G5.TCRT (Supplementary Table 1). Over-expression of MR1*01 in these cells resulted in 7G5.TCRT producing IFNγ and killing the cells at a level equivalent to the reactivity seen with A549, which are heterozygous for MR1*01/MR1*04 (Figure 2a). To confirm the *in vitro* findings and mitigate against cell culture artefacts impacting *in vitro* results, two different *in vivo* models were established in immune-compromised mice. One utilized NALM-6-luciferase, an aggressing leukemic model naturally heterozygous for MR1*04. The second model utilized the melanoma xenograft A375 (MR1*01 homozygous) engineered to express supra physiologic levels of MR1*01. Both models demonstrated robust delay to tumor progression compared to control untransduced donor T cells (Supplementary Figure 3). Although activated by supra-physiologic levels of MR1*01, we investigated if the R9H substitution that differentiates MR1*01/MR1*02 and MR1*04 could further increase 7G5 activation. Previously generated C1R overexpressing MR1^R9H^ (C1R.MR1^R9H^)^12^, were sorted for low and high expression (C1R.MR1^R9Hlo^ and C1R.MR1^R9Hhi^) and used to activate Jurkat.β2m^ko^ expressing the 7G5 TCR (Jurkat. β2m^ko^.7G5 cells). Despite overall lower MR1 surface expression than C1R.MR1, C1R.MR1^R9Hlo^ and C1R.MR1^R9Hhi^ activated Jurkat.β2m^ko^.7G5 to a greater extent than C1R.MR1, as measured by CD69 surface expression (Figure 2b, Supplementary figures 4 and 5).

### 7G5 reactivity is similar to other MR1-restricted TCRs for allele and ligand discrimination

MR1-restricted T cells show diverse TCR usage, exhibit varying distribution, cell reactivity, and ligand preferences. We investigated whether the preferential MR1*04 reactivity seen with the 7G5.TCRT was shared by TCRs derived from additional T cell clones (759S, A4 and C1; called “7G5-like” here). The 759S TCR was derived from the T cell clone MC.27.759S as previously described^25^, and TCRs A4 and C1 were derived from T cell clones isolated by similar methodology to the identification of the MC.27.759S clone. We also compared reactivity of 7G5.TCRT with MR1-reactive TCRs derived from T cell clones AVA34, DGB129 and TCA5A87, reported elsewhere^32^, raised with different methodology, and termed “MR1T”. Jurkat cells CRISPR-engineered for deletion of endogenous TCR and β2M (to prevent aberrant TCR chain pairing and recognition of MR1 *in trans* on adjacent Jurkat cells) were transduced with the described TCRs, and co-cultured with cancer cell lines expressing MR1*01, MR1*02, MR1*04, or C1R cells engineered to express supra-physiologic MR1*01. Jurkat reactivity as measured by CD69 upregulation, fell into 3 main categories (Figure 3a). Jurkat cells expressing 7G5, the 7G5-like TCR 759S, or the MR1T TCR TC5A87 displayed a similar pattern of reactivity against cancer cell lines, exhibiting elevated reactivity to MR1*04 cells and limited reactivity to MR1*01 and MR1*02 lines. Jurkats expressing one of the remaining 7G5-like TCRs, A4 or C1, reacted to MR1*04 but also showed greater CD69 upregulation to some cell lines expressing MR1*01 and MR1*02, compared to 7G5. Jurkats expressing the MR1T cell derived TCRs AVA34 or DGB129 showed no apparent reactivity to cancer cell lines with physiologic MR1 expression. All the TCR engineered Jurkats were robustly reactive to the positive control cells C1R expressing supra-physiological levels of MR1*01.

**Figure 3.**
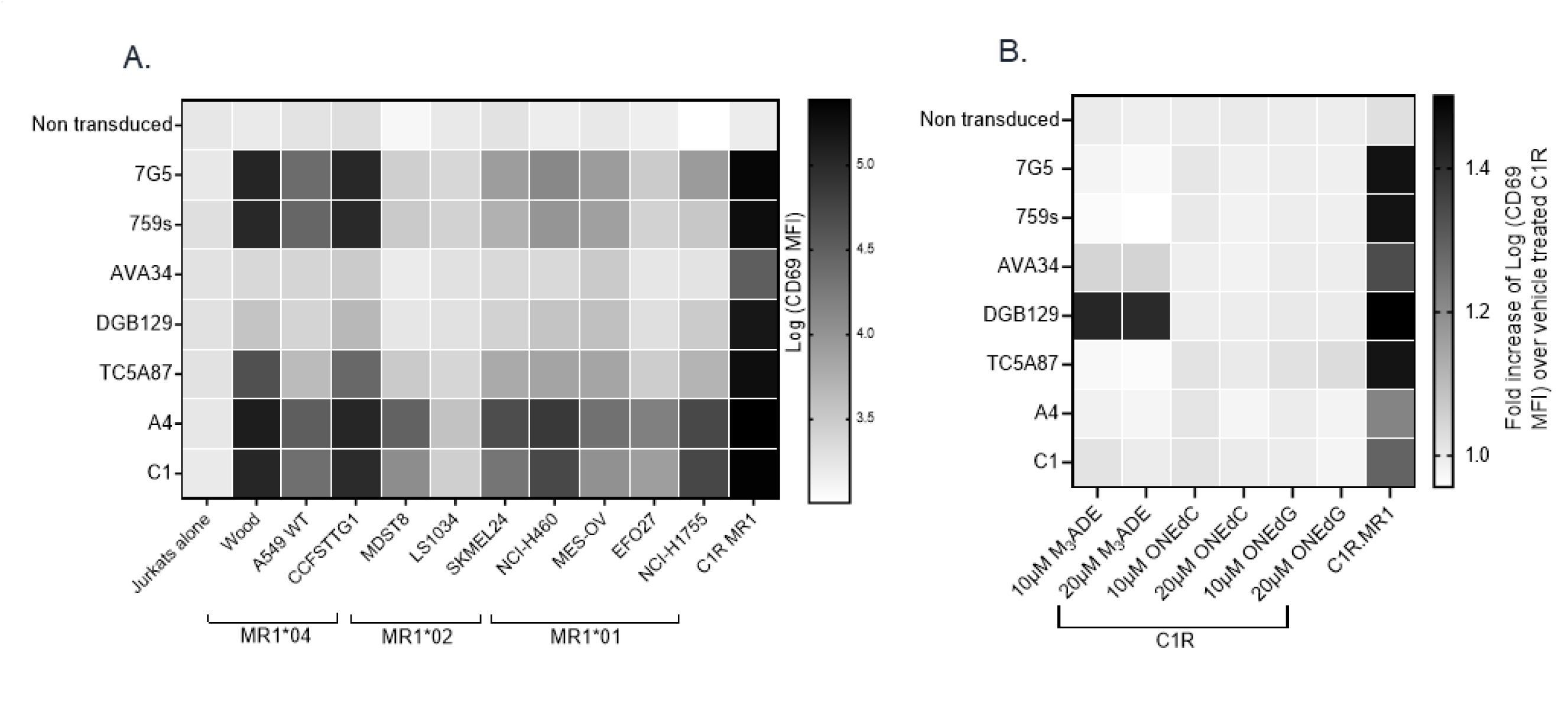
Comparison of MR1 allele restriction and ligand reactivity of 7G5-TCRT with other MR1-reactive TCRs. **3a** Heatmap showing log median fluorescence intensity values of CD69 surface levels on Jurkat cells expressing one of seven T cell receptors (Y axis) after 24-hour incubation with MR1*01, MR1*02 or MR1*04-expressing cancer cell lines (X axis). Jurkat cells were incubated alone or with C1R cells overexpressing MR1*01 as negative and positive controls respectively. Geometric mean (n=3) values are plotted. Data are representative of two experimental repeats. **3b** Heatmap showing log median fluorescence intensity values of CD69 surface levels on Jurkat cells expressing one of seven T cell receptors (Y axis) after 24-hour incubation with C1R cells in the presence of the indicated ligand concentrations (X axis). C1R cells were pre-incubated with media or vehicle alone as negative controls, and Jurkat cells were incubated alone or with C1R cells overexpressing MR1*01 as negative and positive controls respectively. Geometric mean (n=3) values are plotted. Data are representative of two experimental repeats.

Collectively these data point to the 7G5-like TCRs and possibly TC5A87 having activity restricted to MR1*04-expressing cell lines in the endogenous MR1 setting.

Some putative metabolite ligands specifically recognized by MR1T have been reported^27^ . We asked whether 7G5-like TCRs could recognize these ligands (3*Z*,5*E*)-6-((9*H*-purin-6-yl) amino) hexa-1,3,5-triene-1,1,3-tricarbaldehyde (“M3ADE”), and 6-((2*R*,4*S*,5*R*)-4-hydroxy-5-(hydroxymethyl)tetrahydrofuran-2-yl)-3-(2-oxoheptyl)-1,8a-dihydroimidazo[1,2-*c*]pyrimidin-5(6*H*)- one (“ONEdC”) presented on C1R cells, which naturally express MR1*01. Using the same Jurkat system, we showed that the TCRs derived from MR1T lines DGB129 and AVA34 recognized C1R targets incubated with M3ADE as reported previously^33^. Pre-incubation of C1R cells with M3ADE or ONEdC did not render C1R cells sensitive to the remaining TCRs, suggesting these ligands are not responsible for 7G5.TCRT, 7G5-like TCRs, or the MR1T TC5A87 liganded MR1 recognition (Figure 3b).

### The 7G5.TCRT does not recognize unliganded MR1

As the 7G5 TCR, in the context of the MC.7.G5 clone, appears to depend on the key lysine residue present in the ligand-binding groove at position 43 of the mature MR1 protein (K43) for its activation^25^ it is likely that this recognition is ligand-dependent. The lysine at position 43 forms a Schiff base with certain metabolites, anchoring them in the MR1 binding cleft^1, 2, 14^. K43 resides deep in the A’ pocket of the MR1 binding groove and is unlikely to be easily accessible to TCRs. We tested the possibility that the 7G5 TCR recognizes MR1*04 in a ligand-agnostic or ligand-independent fashion. We reasoned that if 7G5 binds to MR1*04 in a ligand agnostic fashion, then saturating the surface MR1 of cancer lines expressing MR1*04 with the bacterial ligand Acetyl-6-formylpterin (Ac- 6-FP) should not block recognition. Ac-6-FP was previously shown to block activity of the MC.7.G5 T cell clone^25^. Howson et al demonstrated that Ac-6-FP is presented in the context of MR1 with a R9H substitution, however the MAIT stimulating ligand 5-OP-RU was not presented by MR1 R9H^12^. For MR1*04 cell lines (Wood, Throne, and A549), Ac-6-FP abrogated the response of 7G5.TCRT as measured by IFNγ production (Figure 4). Thus, these data would support the idea that the 7G5 TCR in the context of TCRT does not recognize MR1 in an unliganded fashion.

**Figure 4.**
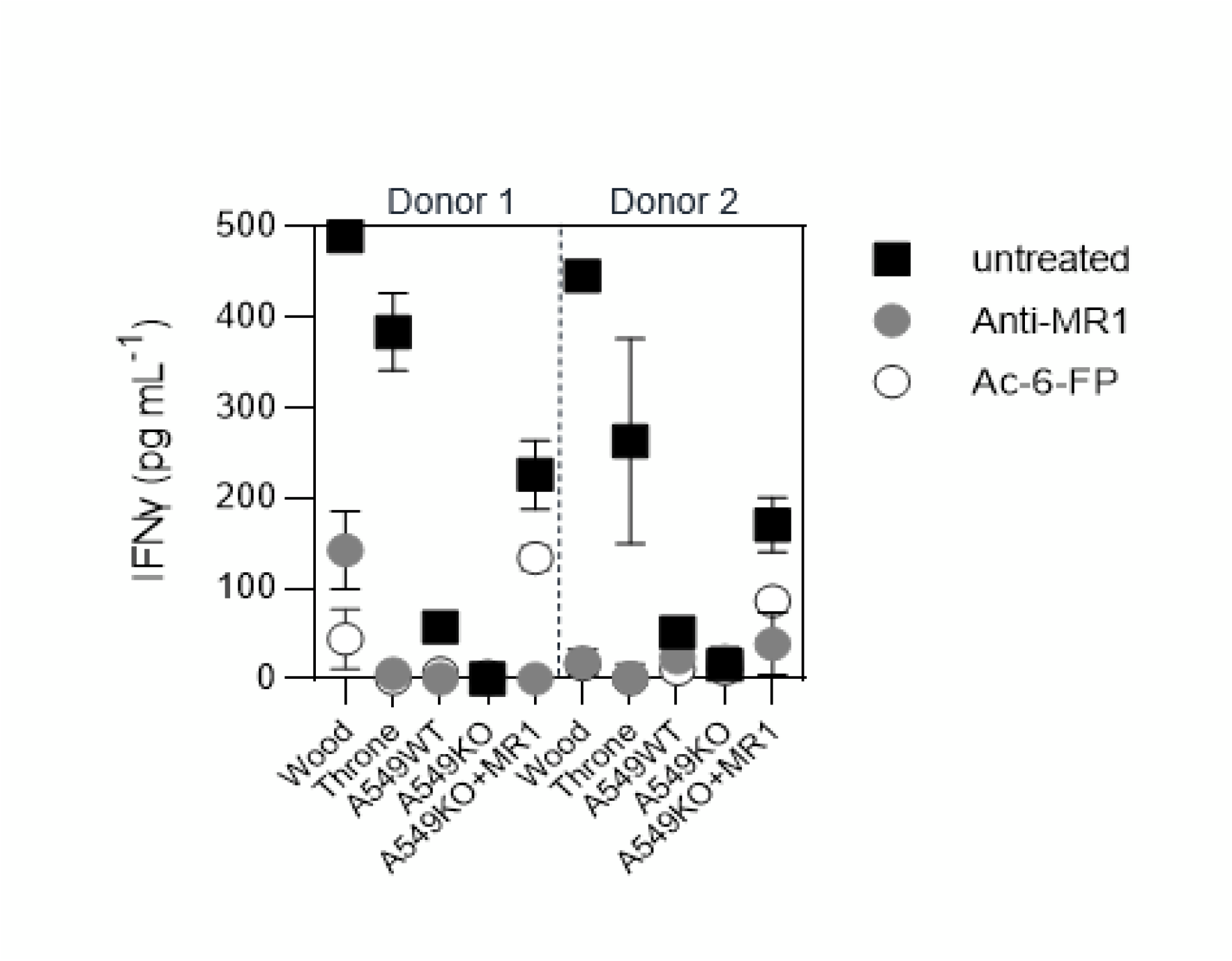
7G5-TCRT does not recognize MR1 in an un-liganded mode. Reactivity of 7G5.TCRT derived from two donors to cancer lines expressing MR1*04, or A549.MR1KO negative control cells. in the presence or absence of Ac-6-FP or blocking anti-MR1 antibody. Was measured by ELISA at 18 hours following co-culture of 20,000 target cells and 60,000 7G5.TCRT, E:T ratio 3:1

### T cells transduced with 7G5, and 7G5-like T cell receptors are activated by MR1*04 expressing healthy non-cancer cells

Given the allele frequency of MR1*04 heterozygotes is estimated to be approximately 1 in 100 in European Caucasians^9^ we reasoned that previous characterization of the MC.7.G5 T cell clone’s ability to be activated in the presence of healthy cells would likely have been performed on cells derived from MR1*01 or *02 allele-expressing donors^25^. We next tested two alternative hypotheses; either 7G5.TCRT recognizes a cancer-specific/enriched ligand restricted by MR1*04 or binds preferentially to a ligand presented by all cells expressing the MR1*04 allele. We sequenced the MR1 locus of ∼200 healthy blood research donors for MR1 allele identities. We identified four donors who were heterozygous for MR1*01/*04 (example plots in Figure 5a). To conduct reactivity assays of 7G5.TCRT against healthy, benign blood cells we isolated monocytes and B cells from three MR1*01/*04 donors as cell types that express the highest cell surface levels of MR1 in blood. 7G5.TCRT did not produce IFNγ or granzyme B in response to monocytes and B cells from MR1*01 homozygous donors. In contrast, significant amounts of both IFNγ and granzyme B were produced in co-cultures with MR1*01/*04 heterozygous donor-derived B cells and monocytes, suggesting that 7G5 TCRT recognize a ligand(s) that is present both in tumor cells and normal cells expressing MR1*04 (Figure 5b). Finally, we tested whether the 7G5-like TCRT (A4.TCRT and C1.TCRT) had similar cancer and normal cell reactivity to 7G5.TCRT. The three TCRT react to cancer cells heterozygous for MR1*04, and only marginally or not at all to cells which express only MR1*01 or are heterozygous for MR1*01/*02 (Figure 5c). Again, this reactivity was not cancer specific, with A4.TCRT and C1.TCRT producing IFNγ and granzyme B in response to B cells and monocytes from MR1*04 heterozygous donors and not in response to MR1*01/02 donors (Figure 5d).

**Figure 5.**
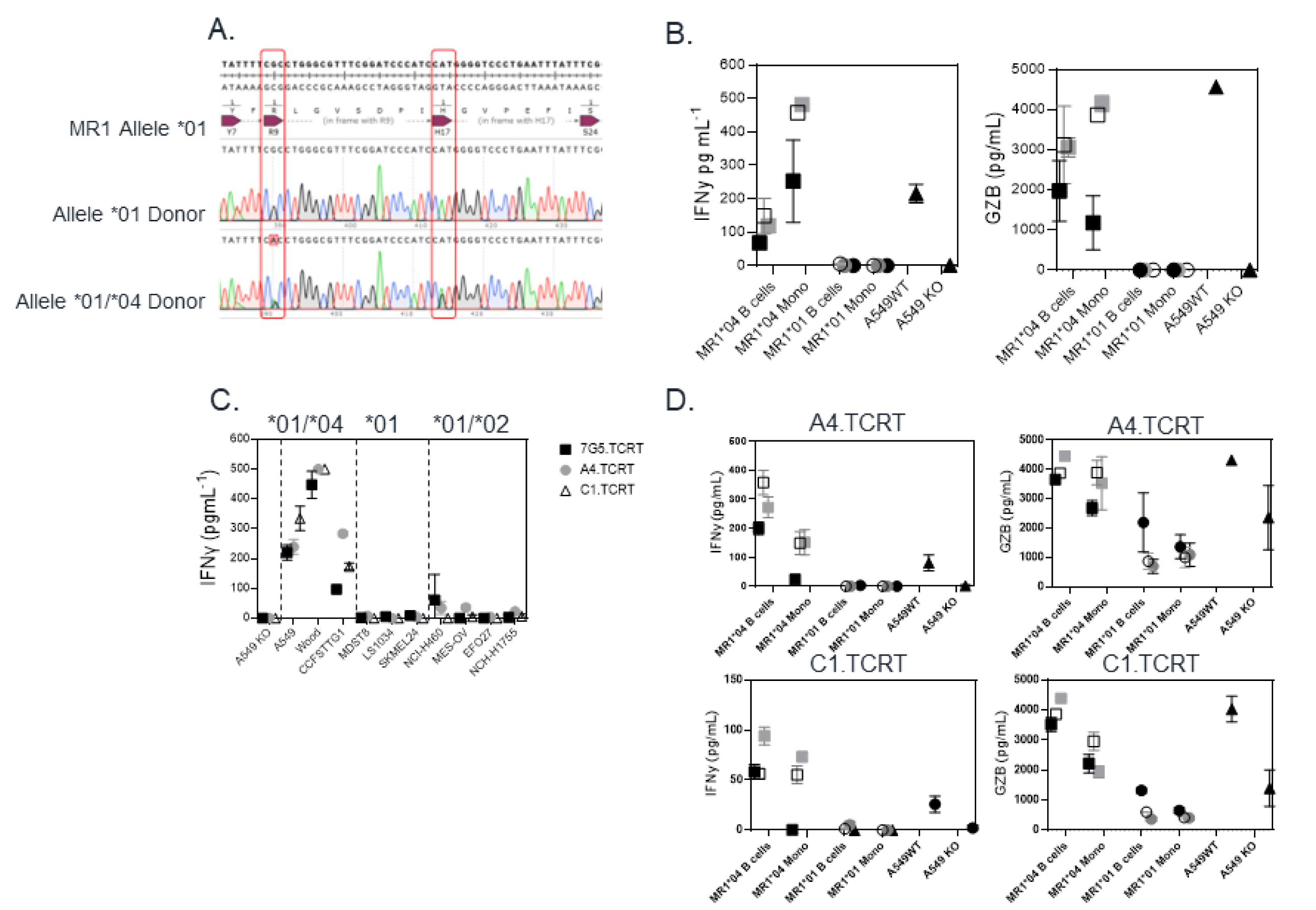
T cells transduced with 7G5 and 7G5-like TCRs are activated by MR1*04 expressing healthy cells. **5a.** Sanger sequencing of the MR1 locus of 2 of 200 blood donors analyzed. The MR1*01 allele and MR1*04 allele differ by 2 nucleotide substitutions leading to R9H/H17R substitutions in the MR1*04 allele. The lower row shows a donor heterozygous for MR1*01/*04, note the mis-calling of the mixed nucleotides at both positions. **5b.** Reactivity of one representative donor of three 7G5.TCRT donors to monocytes and B cells derived from PBMC from three MR1*01/*04 heterozygous blood donors, and three MR1*01 homozygous donors. IFNγ and Granzyme B were measured by ELISA at 48 hours following co-culture of 0,000 PBMC derived cells and 200,000 7G5.TCRT, E:T ratio 3:1.Grey, black, and white squares represent monocytes and B cells from three MR1*01/*04 donors. Grey, black and white circles represent monocytes and B cells from three MR1*01 donors. Black triangles represent the positive and negative controls A549 and A549.MR1KO respectively. **5c.** Reactivity of the 7G5.TCRT and TCRT expressing the 7G5-like TCRs A4 and C1 to cancer cell lines homozygous for MR1*01 or heterozygous for MR1*01 and MR1*02 or MR1*04. Reactivity was measured wit IFNγ ELISA at 48 hours following co-culture of 20,000 cancer cells and 60,000 7G5.TCRT, E:T ratio 3:1 and data shown are representative of 2 biological repeats with two separate donors for TCRT. **5d.** A4.TCRT and C1.TCRT reactivity of one representative of three TCRT donors to monocytes and B cells derived from PBMC from three MR1*01/*04 heterozygous blood donors, and three MR1*01 homozygous donors. IFNγ and Granzyme B were measured by ELISA at 48 hours following co-culture of 40,000 PBMC derived cells and 200,000 7G5.TCRT, E:T ratio :1. Grey, black, and white squares represent monocytes and B cells from three MR1*01/*04 donors. Grey, black, and white circles represent monocytes and B cells from three MR1*01 donors.

## Discussion

In recent years MR1 has held great promise as a target for TCR-mediated tumor immunotherapy. This is largely because it was considered to be monomorphic, invariant across all populations. MR1- restricted T cell clones have been identified which could kill multiple cancer lines without recognizing healthy, non-cancer cells raising the potential of translating TCR-based, MR1-restricted therapy for cancer patients^24–26^. The discovery of rare T cell clones capable of recognizing potential “pan-cancer” MR1-restricted ligands has led to intense efforts to learn more about the biology of MR1 and MR1- restricted T cells in cancer. Recently it has been shown that MR1 has at least 6 alleles. The substitution R9H associated with the MR1*04 allele potentially skews the ligands that are able to be presented by MR1*04 protein, as inferred by the observation that it is unable to present 5-OP-RU^9, 12^. We set out to test the idea that therapeutic TCRT cells could be generated, expressing TCRs from T cell clones able to kill cancer cells from multiple tumor types while leaving benign cells untouched^24, 25^.

In this study we have shown that the 7G5 TCR derived from the MC.7.G5 T cell clone, and other MR1-restricted TCRs with similar properties (e.g., tumor cell recognition and MR1 K43-dependence), robustly re-direct T cells to kill cancer cells in the TCRT format both *in vitro* and *in vivo*. Whilst validating the 7G5.TCRT we observed that robust activity was not pan-cancer in nature, being restricted to a minority of cancer cell lines. Efforts to identify a biomarker for reactivity of 7G5.TCRT led to the discovery that one rare (approximately 1:100) allele of MR1, MR1*04, predicted cancer cell line reactivity. Ultimately, this led to the determination that 7G5.TCRT was not cancer specific, but reactive to normal B-cells and monocytes heterozygous for MR1*04.

To our knowledge this is the first demonstration of an MR1-restricted TCR which has MR1 allomorph specificity. Although the ligands for 7G5.TCRT and related TCRs have not yet been identified, we demonstrated that ligand is likely required to be bound to the MR1 binding groove for 7G5.TCRT reactivity. We have also shown that 7G5.TCRT TCRs do not recognize previously described MR1T ligands at physiological levels of MR1*01 expression in cancer cell lines. A recent study^33^ demonstrated that conventional MAIT cells from healthy donors have the propensity to be promiscuous in their recognition. Such cells express markers of conventional MAIT cells and become activated in response to multiple ligands presented by MR1*01 over-expressed on cancer lines. These cells can react to healthy monocyte derived dendritic cells, but the ligands recognized in this case remain to be identified.

The MR1*04 allele differs from MR1*01 by two nucleotide variants, resulting in an R9H substitution in the ligand binding cleft and an H17R substitution in the α-1 domain outside the binding cleft. Howson et al identified a homozygous MR1 mutation MR1^R9H/R9H^ in a patient with a primary immunodeficiency which was characterized by tattoo-associated human papilloma virus–positive (HPV+) warts^12^. The patient displayed selective loss of all MR1 restricted MAIT cells due to this mutation, which caused structural changes to the ligand-binding pocket of MR1. This mutation accommodates binding of Ac-6-FP but precludes binding of the stimulatory riboflavin-based MAIT ligand 5-(2-oxopropylideneamino)-6-D-ribitylaminouracil (5-OP-RU), and subsequent ability of antigen presenting cells to activate in an MR1-restricted manner. Lack of binding to 5-OP-RU results in loss of ability to upregulate MR1^R9H^ in response to this ligand. The authors showed the patient had an expanded Vγ9/Vδ2+T cell population, which may have arisen to compensate for the loss of circulating MAIT cells. The R9 residue of MR1 is a known MAIT TCR contact, as shown in the crystal structure data of MR1 binding the drug diclofenac^21^, and as such may be involved in binding to other ligands. Our studies show that 7G5.TCRT and 7G5-like TCRs can react to targets expressing supra- physiologic MR1*01 and MR1*04 suggesting that either multiple ligands can be recognized by these TCRs, or the same ligand is recognized but is presented preferentially with the histidine at position 9 in the binding cleft of MR1*04. The significance of MR1*02 for MR1T and MAIT cell function is unknown. MR1*02 does not appear to be recognized at physiological levels by 7G5.TCRT. Approximately 25% of the population carry the MR1*02 allele which incorporates the H17R substitution^9^. The location of arginine 17 as revealed by the MR1 crystal structure^1^ is distant to the binding cleft so is unlikely to be involved in antigen binding or TCR binding. It is possible that arginine 17 plays a role in MR1 trafficking or antigen loading, which might have applied evolutionary selection pressure to retain this allele.

The 7G5 TCR appears to be exquisitely sensitive to MR1*04 when the TCR is expressed either in Jurkat cells or primary T cells. A549 cells are one of the most stimulatory lines for 7G5.TCRT, but MR1 (MR1*01 and/or MR1*04) is barely detectable on their surface by flow cytometry. This does not appear to be a technical artifact as the antibodies most commonly used to detect MR1 by flow cytometry 8F2.F9^12^ and 26.5, used in this study, both recognize MR1*01 and MR1*04 allomorphs^12^. This reactivity contrasts with MR1T TCRs such as DGB129^26^ and TC5A87^24^ which in these studies appear to rely on targets that have been engineered to over-express MR1^26^. This may simply reflect the methodology used to isolate such cells, with targets either over-expressing MR1 or not. Our data demonstrate that 7G5.TCRT have up to 110-fold greater reactivity to MR1*04 but can still react to over-expressed MR1*01. As such, it is important for future studies to consider the potential biological implications of observations made with physiological versus over-expressed MR1 protein.

We do not yet know the ligand(s) bound to MR1 required for 7G5.TCRT activity. However, we do know that the putative MR1T ligands ONEdc or M3ADE do not activate 7G5.TCRT or TCRs identified using comparable methods. This is perhaps unsurprising as previous work has shown that, unlike the MR1T cells, the 7G5 TCR relies on the MR1 binding groove residue K43 for ligand recognition^25^, likely through a Schiff base formed with the ligand. For its role in tumor surveillance, our data challenge the current hypothesis that the ligands seen by MR1T, and 7G5-like T cells are cancer specific metabolites^24, 25^. Our findings suggest that the ligand(s) of 7G5 are common to normal and transformed cells, challenging this recent dogma, or the TCR may be recognizing more than one ligand.

The original MC.7.G5 T cell clone is heterozygous for MR1*01/*02 (Supplementary Figure 6) and therefore would not have been negatively selected on MR1*04 in the thymus. The patient with the reported MR1^R9H/R9H^ genotype had no detectable MR1-restricted T cells^12^ . This suggests that being homozygous for this polymorphism, and perhaps for MR1*04, may not be compatible with thymic positive selection of MR1-restricted T cells. It is clear from our data that healthy individuals lacking MR1*04, such as the donor of MC.7.G5, can harbor T cells which are not negatively selected in the thymus and can recognize a ligand or ligands derived from the normal metabolome or proteome and restricted by MR1*04. This finding has potential implications for transplantation. It is conceivable that allogeneic transplantation from donors lacking MR1*04 to those expressing this allele could pose a risk of graft versus host disease (GVHD). Conversely, transplantation of organs from MR1*04 donors to negative recipients may represent a higher risk of allogeneic rejection.

The promise of MR1-restricted TCR-based therapy for cancer has arisen through the discovery of T cells from human donors that appear to be restricted to MR1-ligand targets preferentially found in cancer. These data demonstrate that for TCRs derived from T cell clones such as MC.7.G5, specificity appears to be restricted to a relatively rare MR1 allele bearing two SNVs and found in approximately 1% of the human population. This observation is important for three reasons. One, it highlights the need for deeper understanding of non-MAIT, MR1-restricted TCRs prior to clinical translation; two, it raises the importance of deciphering the ligands driving reactivity in cancer and infection and finally, it raises the specter that MR1 allele variants warrant understanding in the context of allotransplantation.

## Methods

### Cell lines and cell culture

Cancer cell lines were cultured according to manufacturer’s instructions.

### Blood derived cells

Whole blood was sourced from research donors via Cambridge Bioscience, under local ethical review panel guidelines. PBMCs were isolated from whole blood using lymphoprep and leukosep tubes according to manufacturer’s instructions. PBMCs were used as starting material to isolate B cells (CD19+ microbeads, Miltenyi Biotec) and monocytes (CD14+ microbeads, Miltenyi Biotec) following manufacturers’ protocols.

### ELISA assays

All co-culture assays were carried out in RPMI containing 10% fetal calf serum. Target cells were plated in flat bottomed 96 well plates, T cells were thawed and rested for 2 hours prior to plating. Target cell numbers, T cell numbers, effector to target ratios can be found in relevant figure legends. For assays comparing TCR from MC.7.G5 to TCRs in clinical development, effector T cells were normalized for percentage of specific TCR positive cells by addition of appropriate numbers of non-transduced T cells from the same donor, to allow for comparison between TCRs. Target and T cell co-cultures were incubated at 37°C in 5% CO_2_ co-culture durations can be found in relevant figure legends. Supernatants were collected from effector-target cell co-cultures and analyzed for IFNy using either ELISA MAX Deluxe Human IFNγ kit (BioLegend) or Human IFNγ DuoSet ELISA (R&D Systems) and for granzyme B using human Granzyme B DuoSet ELISA (R&D Systems). All ELISAs were performed as per manufacturer’s instructions. ELISAs were read on a Mini ELISA plate reader from Biolegend.

### Ac-6FP and anti-MR1 cell treatment

Target cells were plated at 20,000 cells per well in a flat bottom 96 well plate and allowed to attach for 2 hours. Following this, cells were pre-incubated with 10μg/mL anti-MR1 clone 26.5 (BioLegend) or 100μM Ac-6-FP for 4 hours before adding 60,000 T cells and incubating for an additional 18 hours at 37°C in 5% CO_2._

### Flow cytometry

Adherent cells were harvested from plates using TrypLE express (Thermo Fisher Scientific) and transferred to a 96-well round bottom plate for staining. Cells were stained with zombie violet viability dye (BioLegend) in PBS for 10 minutes at 4°C, followed by anti-MR1 APC (clone 26.5, Biolegend) or isotype control (MOPC-173, Biolegend) in FACS buffer for 30 minutes at 4°C. Cells were washed twice, resuspended, and acquired on a Beckman Coulter Cytoflex-S flow cytometer. Data was analyzed using FlowJo software.

### TCR sequences

Genes encoding the full length TCR-α and TCR-β chains linked by a 2A-furin sequence for AVA34 DGB129 and TC5A87 according to the sequences in patent WO2021144475A1, c259 from patent WO2020049496A1, ADP-A2M4 from patent US9976121B2 and IMA203, from patent US20200376031A1, were synthesized and cloned into the vector pSF-LV-EF1a (Oxgene) using the GeneArt service (ThermoFisher Scientific).

### Transfection of HEK293T Lenti-X cells and virus concentration

HEK293T Lenti-X cells were transfected using PEIpro (Polyplus) as per manufacturer’s instructions with a lentivector encoding the appropriate T cell receptor or MR1 construct alongside pREV.Kan, pGagPol.Kan, and pVSVG.Kan packaging vectors (Aldevron). After 72 hours, virus-containing cell supernatants were concentrated using Amicon Ultra-15 Centrifugal Filter Units (Merck Millipore) and exchanged into TexMACS media (Miltenyi Biotec).

### Cell line transduction with MR1

Cell lines to be transduced were thawed and rested overnight in RPMI supplemented with 10% fetal calf serum at 37°C and 5% CO_2_. After resting, cells were harvested, counted, and plated at 300,000 cells per well in a 6-well plate. Cells were then transduced to express B2M-MR1, using concentrated supernatant from transfected HEK293T, as described above, and polybrene (dilution 1:1000). Efficiency was analyzed 6-7 days post-transduction using flow cytometry.

### Lentiviral transduction of primary T cells and Jurkat cells

Jurkat E6.1 TCRαβ-β2M- CD8+ NF-κB:CFP NFAT:eGFP AP-1:mCherry cells (a kind gift from Peter Steinberger, Medical University of Vienna) were transduced to express a TCR using concentrated supernatant from transfected HEK293T cells and Polybrene. Transduction efficiency was analyzed 3- 4 days following transduction using flow cytometry. CD3+ T cells were purified from human peripheral blood Leukopaks (Hemacare) using Cell Therapy Systems CD3/CD28 Dynabeads (Thermo Fisher Scientific). Positively selected CD3+ cells (15 x 10^6^) and Dynabeads in TexMACS 5% human AB serum 20ng/mL IL-2 were seeded into 10M G-Rex systems (Wilson Wolf) and incubated overnight at 37°C 5% CO_2._ CD3+ cells were transduced to express a TCR with or without CD8 or to knockout the TRBC1 and 2 genes with CRISPR Cas9 using gRNAs described by Legut et al^34^ or mock transduced using concentrated supernatant from transfected HEK293T cells and LentiBOOST (Sirion Bio). Following a further 24 hours, fresh TexMACS 5% human AB serum 20ng/mL IL-2 was added to G-Rex systems, where the cells had been transduced with CRISPR Cas9 puromycin was added at a final concentration of 1 ug/ml. Ten days following transduction and expansion, CD3+ cells were harvested, magnetic beads removed, and frozen for storage.

### Generation of the JRT76 β2m^ko^ cell line via CRISPR/Cas9

Jurkat clone 76 (JRT76) cells^35^, a TCRα/β negative derivative of the Jurkat E6.1 cell line (ATCC TIB-152™), were kindly provided by Prof Mirjam H M Heemskerk, Leiden University. Knockout of β2m was performed using the pLentiCRISPR v2 CRISPR/Cas9 system^37^ using a pLentiCRISPR v2 plasmid encoding the β2m-targeting guide RNA (gRNA): GAGTAGCGCGAGCACAGCTA (Genscript). Lentiviral particles were packaged in HEK293T cells (ATCC: CRL-3216™) by co-transfection of pLentiCRISPR v2-β2m-gRNA with pRSV-rev (Addgene Plasmid #12253), pMDLg/pRRE (Addgene Plasmid #12251) & pMD2.G (Plasmid #12259) plasmids which were gifts from Didier Trono, obtained through Addgene. Transfections were performed by complexing plasmids with FuGENE® 6 Transfection Reagent (Promega) in OptiMEM media (ThermoFisher Scientific) before pipetting dropwise onto plated HEK293T cells in D10 culture media: Dulbecco’s Modified Eagle Medium (DMEM) supplemented with 10 % heat inactivated foetal calf serum (FCS), 2 mM GlutaMAX™, 100 units/mL penicillin and 100 µg/mL streptomycin (all ThermoFisher Scientific). Transfections were incubated for ∼16 hrs after which cells were replenished with fresh D10 media. Viral particle containing media was harvested after a further 24 and 48 hrs and pooled.

Target JRT76 cells were transduced by culturing 6.0×10^5^ cells in neat media containing harvested lentivirus. After ∼16 hr incubation with lentivirus, cells were replenished with enriched-R10 media: RPMI supplemented with 10 % heat inactivated FCS, 2 mM GlutaMAX™, 100 units/mL penicillin and 100 µg/mL streptomycin, 0.02 M HEPES, 1 mM nonessential amino acids and 1 mM sodium pyruvate (all ThermoFisher Scientific). After 72 hrs in culture, transduced cells underwent antibiotic selection via addition of 0.5 μg/mL puromycin to culture media for 7 days. Transduction/knockout efficiency was assessed by surface HLA-A,B,C expression; staining cells with W6/32 clone hybridoma supernatant which was detected via a Goat anti-mouse-Cy5 secondary (ThermoFisher Scientific). Subsequent staining was analysed via flow cytometry using a BD® LSR II Flow Cytometer. From the parent transduced line, a subclonal population of JRT76 β2m-knockout cells which exhibited complete absence of HLA-A,B,C expression was achieved via limit dilution cloning.

### Retroviral transduction of JRT76 β2m^ko^ with 7G5 TCR

Jurkat.β2m^ko^.7G5 cells were generated by retroviral transduction of JRT76 β2m^ko^ with p-MIG containing cDNA for the expression of the 7G5 TCR α and β chains separated by a 2A peptide linker for co-expression as described previously^36^ . Cells were sorted for increased CD3 and GFP expression by flow cytometry after staining with anti-CD3-PECy7. Cells were maintained in RF10: RPMI 1640 supplemented with 2 mM MEM nonessential amino acid solution (Gibco), 100 mM HEPES (Gibco), 2 mM L-glutamine (Gibco), Penicillin/Streptomycin (Gibco), 50 µM 2-mercaptoethanol (Sigma-Aldrich) and 10% heat inactivated foetal bovine serum (Sigma-Aldrich).

### Activation assays with Jurkat.β2m^ko^.7G5

Jurkat.β2m^ko^.7G5 cells were co-incubated at a 1:1 ratio with C1R derivatives (10^5^:10^5^) in 96 well U-bottom plate overnight at 37°C, 5% CO_2_ in RF10. Cells were stained for viability (LIVE/DEAD™ fixable Aqua stain, ThermoFisher Scientific) and surface expression of CD3 (CD3 PE-Cy7, clone SK7, BD) and CD69 (CD69 APC, clone L78, BD) in PBS, and fixed with 1% paraformaldehyde in PBS prior to flow cytometry acquisition. Flow cytometry acquisition was performed on an BD LSRII flow cytometer run with BD FACSDiva software (FlowCore, Monash University), and analysed using FlowJo^TM^ software. Cells were gated on FSC-A vs SSC-A, FSC-A vs FSC-H, GFP vs Live/Dead, GFP vs CD3 and the % CD69^hi^ cells extracted (Supplementary Figure 5). In parallel, on the same day, C1R derivatives were stained for MR1 expression using 8F2.F9 hybridoma supernatant, followed by anti-mouse IgG-PE (Goat F(ab’) 2 Anti-mouse IgG (H+L) human ads-PE, Southern Biotech). Cells were gated FSC-A vs SSC-A, FSC-A vs FSC-H and the Median Fluorescence Intensity for PE extracted (Supplementary Figure 4). Data were plotted with GraphPad Prism (GraphPad Software, San Diego, California USA).

### Viral copy number analysis

Virus Copy Number (VCN) was calculated from the concentrations of a viral gene (Psi) and a reference gene (RPP30) measured from the same sample. Genomic DNA was extracted using a Maxwell RSC Instrument (Promega, USA) following the manufacturer’s protocol. Extracted DNA was digested using an EcoRI HF Restriction Enzyme (NEB, USA). The concentrations of Psi and RPP30 were measured in a QIAcuity Digital PCR instrument (Qiagen, Germany) following the manufacturer’s protocol. The following controls were included in each experiment: positive controls with known VCN and no-template controls. The PCR program was: 95°C for 2 min initial heat inactivation, followed by 40 cycles of 15 s denaturation at 95°C and 30 s combined annealing/extension at 60°C. Primers, probes, positive controls, and buffers were purchased from Integrated DNA Technologies (IDT, USA) unless otherwise stated.

Forward primer, Psi: CAG GAC TCG GCT TGC TGA AG

Reserve primer, Psi: GCA CCC ATC TCT CTC CTT CTA GC

Probe, Psi: /56-FAM/TT TTG GCG T/ZEN/A CTC ACC AG/3IABkFQ/

Forward primer, RPP30: AGTGACTGATGCAGGACATTAC

Reserve primer, RPP30: CAGGGCAGAAGAGGCAAATA

Probe, RPP30: /5HEX/AC GCT GTG T/ZEN/G TGG ATT TCT CCT GA/3IABkFQ/

### Cytotoxicity Assays

All cytotoxicity assays were performed using the xCELLigence RTCA MP analyzer and 96 well PET E-plates (Agilent Technologies). Assays were carried out in RPMI containing 10% fetal calf serum. Prior to target cell seeding background measures were taken using assay media. Target cells were then seeded at a density pre-determined to be optimal for each cell line. Plates were incubated overnight at 37°C 5% CO_2_ on the xCELLigence with cell index being measured every 15 minutes. Effector T cells were thawed and prepared as described for ELISA assays and seeded into E-Plates at a ratio of 5:1 T cells:target cells per well. E-Plates were incubated for a further 48 hours at 37°C 5% CO_2_ in the xCELLigence with cell index being measured every 5 minutes. Using the analyzer software, raw cell index values were converted to percentage cytolysis using full lysis (target cells incubated overnight in E-Plates before addition of Tween-20 to a final concentration of 0.25%) and target cell only controls and normalized to the time point before T cell addition.

### Co-culture of TCR-expressing Jurkat cells and ligand-loaded C1R cells

C1R or C1R cells over-expressing MR1*01 (C1R.MR1, a kind gift from Andrew Sewell, University of Cardiff) were stained with CellTrace Violet (ThermoFisher Scientific) as per the manufacturer’s instructions. Cells were then incubated in 96 well flat bottom plates with M_3_ADE, ONEdC, or ONEdG, a kind gift from Prof Del Besra, University of Birmingham) to a final concentration of 10µM or 20µM in RPMI 10% FBS Penicillin/Streptomycin (R10), vehicle only (0.2% DMSO in R10), or R10 only, for 5 hours at 37°C 5% CO_2_. Jurkat cells (5E4 cells) transduced with one of eight TCRs, or non-transduced, were added to the C1R-containing wells at a 1:1 ratio and incubated for 24 hours at 37°C in 5% CO_2_. C1R only samples were stained with Zombie NIR Fixable Viability Kit (BioLegend). Samples were analyzed on CytoFLEX S (Beckman Coulter). Events were gated on FSC-H vs. SSC-H, FSC-H vs. FSC-W, Zombie NIR negative, and then the Median Fluorescence intensity in the PE channel was extracted. Co-culture samples were stained with Zombie Violet Fixable Viability Kit (BioLegend) followed by anti-CD69 APC antibody (FN50, BioLegend). Samples were analyzed on CytoFLEX S (Beckman Coulter). Events were gated on FSC-H vs. SSC-H, FSC-H vs. FSC-W, Zombie Violet/CellTrace Violet negative, and then the Median Fluorescence intensity in the APC channel was extracted. Flow cytometry data was analyzed using FlowJo. Other data was analyzed using GraphPad Prism.

### Sequencing of MR1 alleles

Genomic DNA from the MC.7.G5 clone was extracted using the DNeasy Blood & Tissue Kit (Qiagen) and from blood by using either DNeasy Blood & Tissue Kit (Qiagen) or Maxwell® RSC Blood DNA Kit. The MR1 locus was amplified from the DNA using PCR with Q5 High-Fidelity DNA Polymerase (New England Biolabs) following the manufacturer’s instructions. To amplify the region of the MR1 gene encoding the R9 and H17 residues, the following primers were used: OP71_forward_genomic_R9Hmut 5’- CACACGTGCACACACAGAGGTG, and OP72_reverse_R9Hmut 5’- GGACAGTCCAGAAGATGCACAGG. PCR products were checked by running on a 1% SyberSAFE agarose gel with 1kb DNA ladder (Invitrogen). PCR products of successful reactions were purified and sequenced by Source BioScience using following primers: OP80_forward_Exon2_seq 5’- GAGCTCTTACGTCCTGTCCAGG, OP81_reverse_Exon2_seq 5’ - GCTACAGCAGGTGCAATTCAGC, OP82_reverse_Exon2_seq 5’- GCGAGGTTCTCTGCCATCC, OP83_forward_Exon2_seq 5’- CAGTGTCACTCGGCAGAAGG, OP69_genomic primer 2F 5’- GAAGAAGGCTGCGTCATCAG and OP72_reverse_R9Hmut 5’- GGACAGTCCAGAAGATGCACAGG. Sequencing data was analyzed using SnapGene software.

### In vivo studies

*In vivo* studies were carried out in compliance with the applicable laws, regulations, and guidelines at Labcorp Drug Development (Ann Arbor, MI) or Epistem Ltd (Manchester, UK). For each study mice were checked regularly for health status and body weight and were sacrificed when pre-determined termination criteria were reached.

NALM6: Groups of (n=5) 6–8-week-old female NSG (NOD.Cg.Prkdc^scid^IL2rg^tmWjl^/SzJ- Jackson Laboratory) mice were injected intravenously (i.v.) on day -3 with 5×10^5^ NALM6 B-cell acute lymphoblastic leukemia cells expressing Luciferase (NALM6-Luc-mCh-Puro) (ATCC/Labcorp). Three days after tumor injection, mice were injected with luciferin intraperitoneally (i.p) and imaged under anesthesia using an IVIS S5 Imaging System, with Bioluminescence Imaging (BLI) data analyzed using Living Image 4.7.1 software (both from Perkin Elmer, Waltham, MA). Using these BLI measurements, mice were distributed into groups ensuring that the mean tumor burden for each group was within 10% of the mean tumor burden for the study population. The mice were untreated or injected i.v. with 1×10^6^, 5×10^6^ or 2×10^7^ human T-cells transduced with a vector to express 7G5.TCR, after knocking out the endogenous TCR. IVIS imaging was then performed three days later, and twice each week thereafter, to follow tumor progression. Imaging data were obtained within 10 min after luciferin injection.

A375-MR1: A375 melanoma cells (ATCC) were transduced with lentiviral particles encoding B2m- MR1*01 to express high levels of MR1*01 at the cell surface. NSG mice were subcutaneously (s.c.) injected with 5×10^6^ A375-MR1 cells. 1 day later the mice were randomized into groups of 8 animals and injected i.v. with vehicle, 2.3×10^7^ non-transduced (NTD), or either 2.3×10^7^ or 1.2×10^7^ T-cells from two different donors (equivalent to 1×10^7^ or 5×10^6^ T-cells expressing 7G5.TCRT). For this experiment the endogenous TCR was not ko’d. Mice were weighed and tumor volume was assessed using calipers three times weekly.

## Acknowledgements

The authors acknowledge the provision of Jurkat triple reporter cells from Dr Peter Steinberger, Medical University of Vienna. The authors acknowledge the Monash University FlowCore Facility for flow cytometry instrumentation and technical support, and the services and facilities of Micromon Genomics at Monash University. The T cell receptor sequences derived from the T cell clones A4, C1, and 759S, were developed as part of a Joint R&D Collaboration between Enara Bio and the University of Cardiff, the authors acknowledge the Cardiff inventors Prof. Andrew Sewell and Dr Garry Dolton. The authors thank David Lewinsohn, Oregon Health and Science University, for advice and helpful conversations about the study. The authors thank Bruce MacLachlan for production and provision of the JRT76 B2M knockout parental line.

## Author contributions

**Conceptualization**- Joseph Dukes, Duncan Howie, Sophie Papa, Luke Williams,

**Methodology**- Suzanne Cole, Terri Cornforth, Eleanor Denham, Patricia Illing, Emily Lam, Angharad Lloyd, Tatiana Lobry, Nathifa Moyo, Jeremy Raynes, Ron Wolchinsky

**Investigation-** Suzanne Cole, Terri Cornforth, Eleanor Denham, Patricia Illing, Emily Lam, Angharad Lloyd, Tatiana Lobry, Nathifa Moyo, Jon Silk, Jeremy Raynes, Katarzyna Ward, Bruce MacLachlan, Julian Vivian

**Supervision- Anthony Purcell, Jamie Rossjohn, Duncan Howie, Luke Williams, Joseph Dukes Resources-** Gurdyal S Besra, Natacha Veerapen, Anthony Purcell, Jamie Rossjohn

**Manuscript writing-**Duncan Howie, Sophie Papa

**Manuscript review and editing-**Duncan Howie, Sophie Papa, Joseph Dukes, Jonathan Silk, Jérôme Le Nours, Anthony Purcell

## Supplementary Figures

**Supplementary Figure 1.**
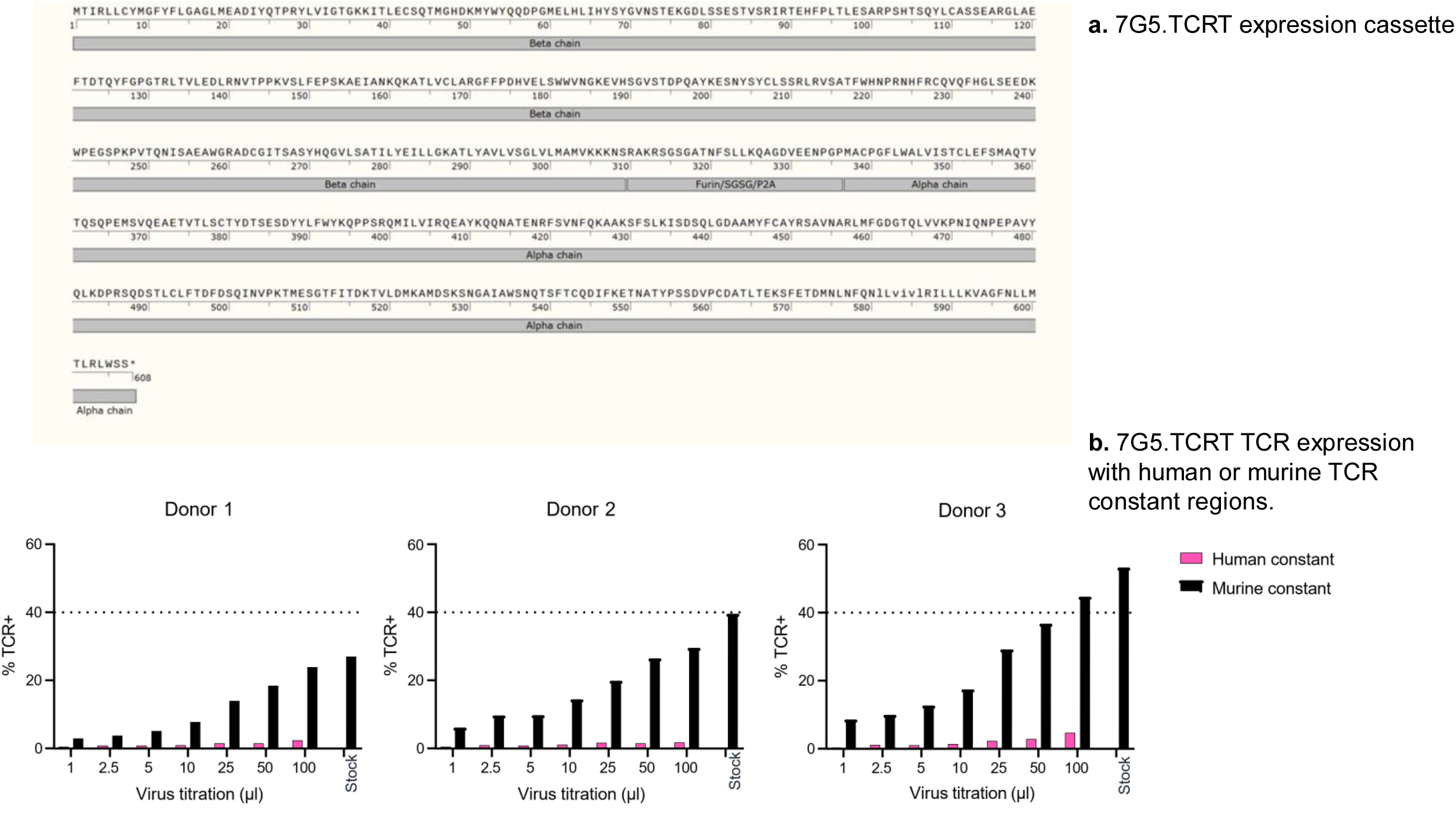
7G5 expression cassette sequence **a.** Map of the 7G5 TCR expression cassette used in this study. Constant alpha and beta regions were murine and hydrophobic residues were introduced into the transmembrane region of the alpha chain as described in the text and denoted in lowercase on the protein sequence **b.** Comparison of 7G5.TCRT manufactured with murine or human TCR constant regions. Concentrated lentivirus was titrated using three T cell donors and 7G5 TCR expression, measured by TRBV25 expression (normal expression on only 1% of peripheral T cells) was measured by flow cytometry.

**Supplementary Table 1.**
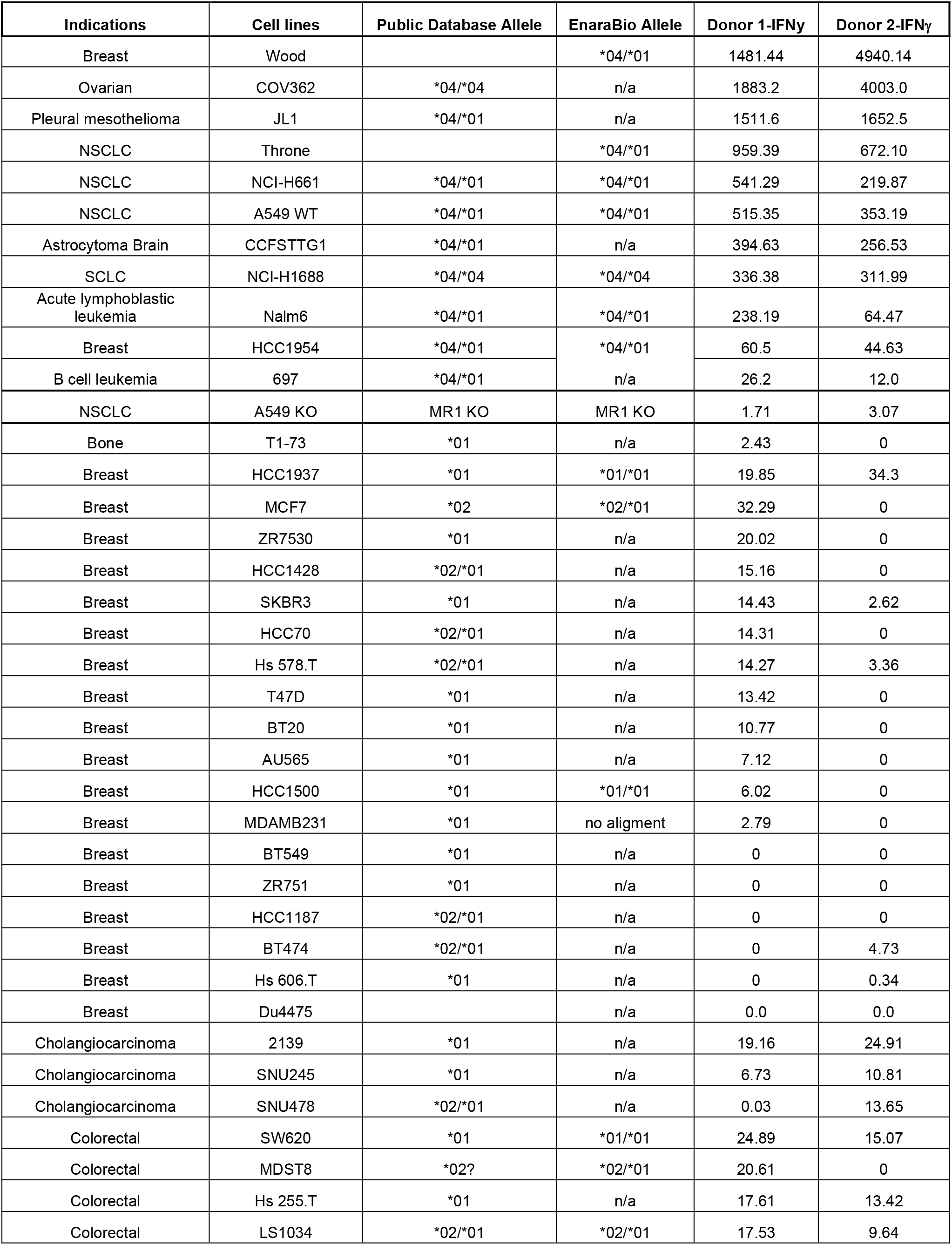
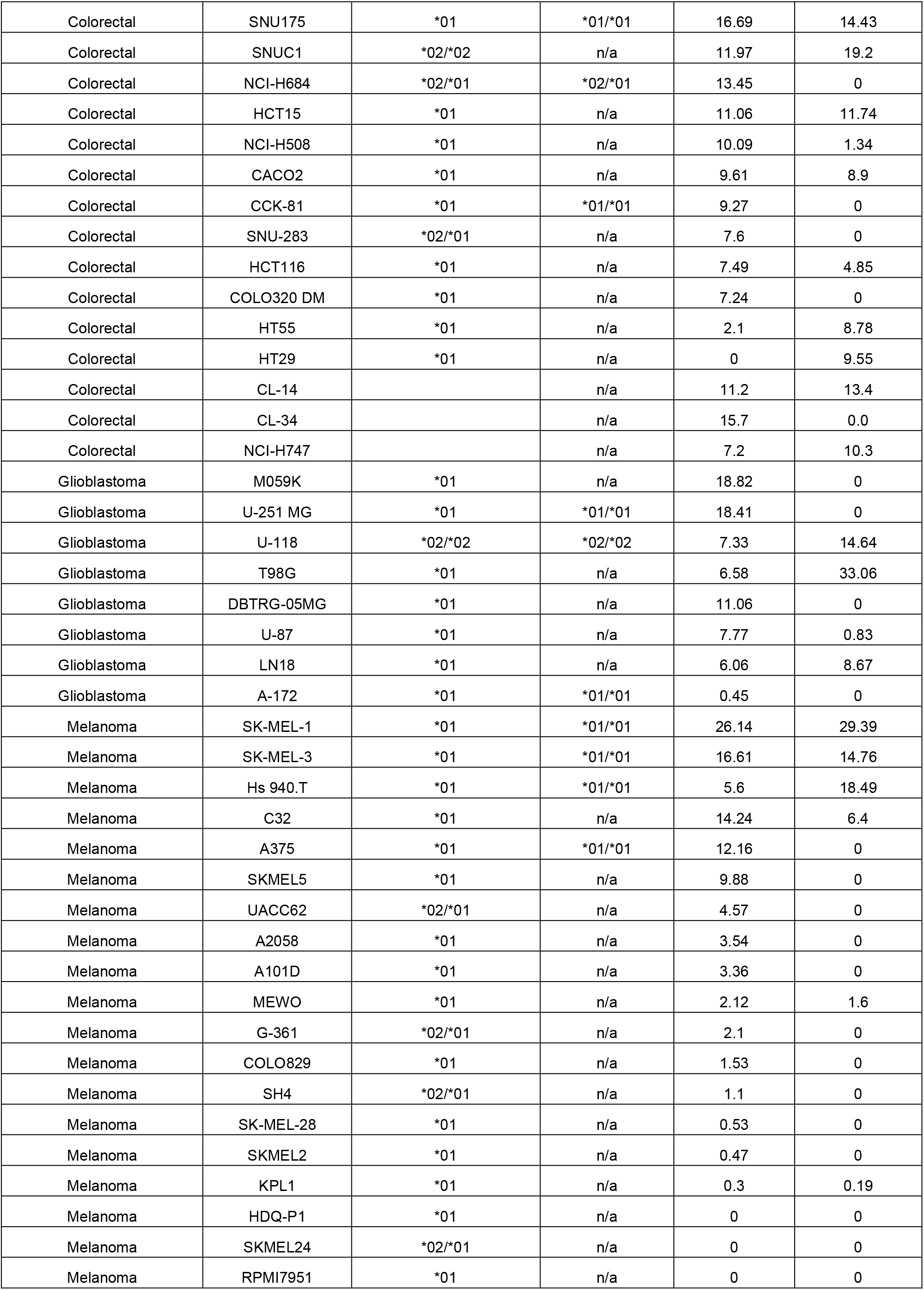
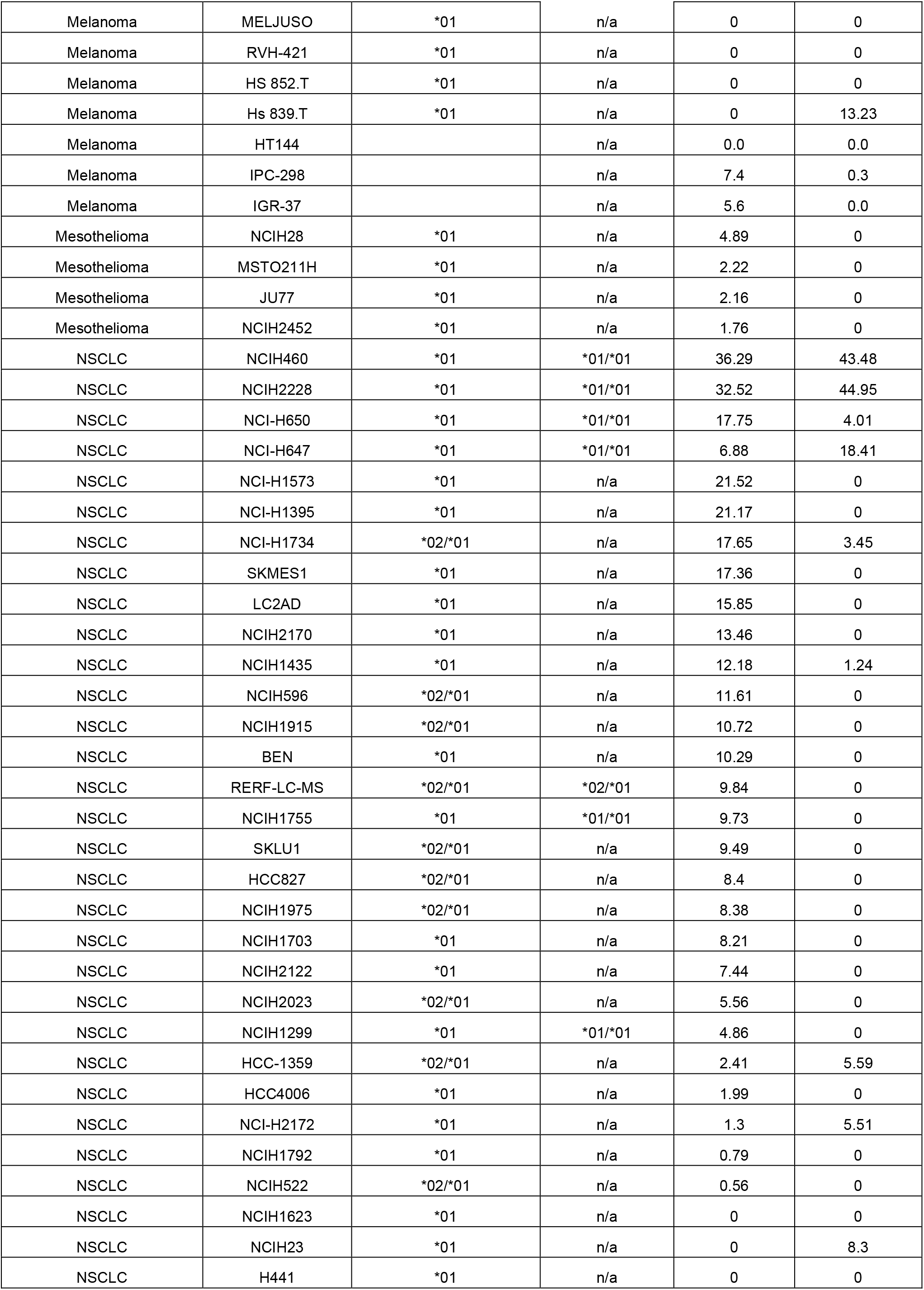
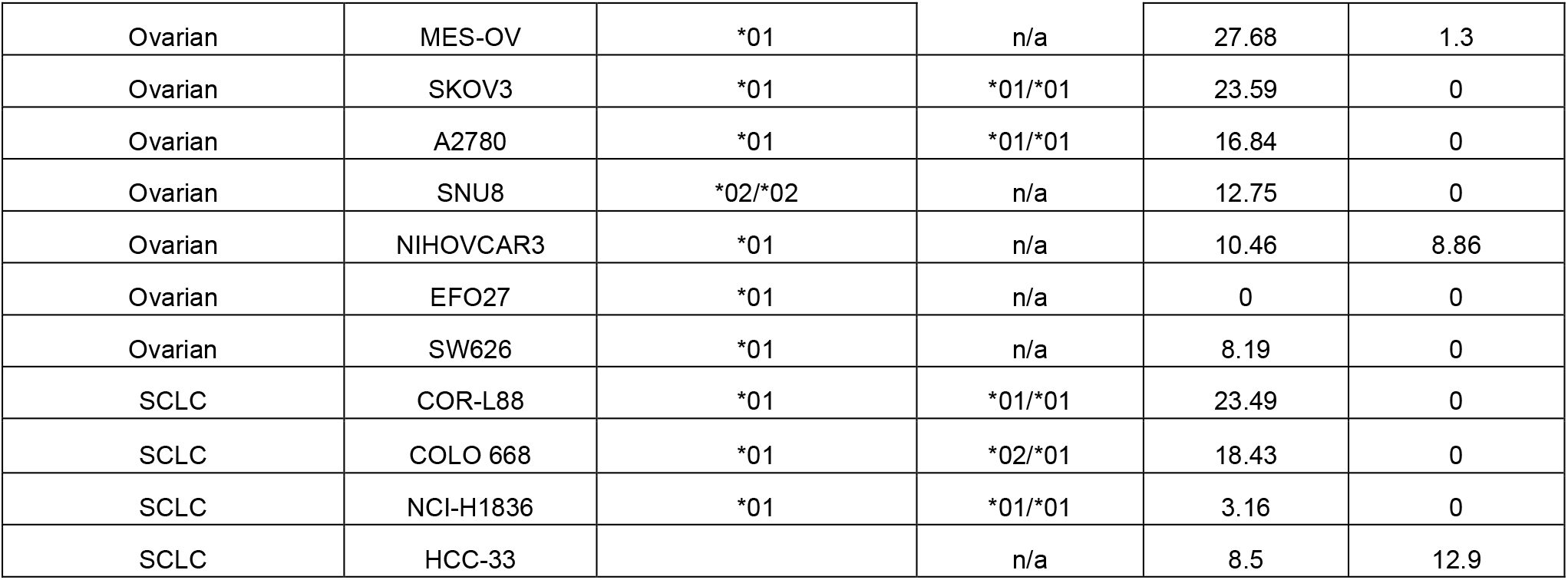
Kill data on 140 cancer cell lines, IFNγ and MR1 allele status. Table showing 7G5.TCRT reactivity as measured by IFNγ ELISA to 135 cancer cell lines, including the MR1 allele expression of the lines. MR1 allele expression was derived from both public databases (CCLE and Crown Bioscience) and from internal PCR-sequences of DNA from these lines. Reactivity data from two separate T cell donors is shown. The table shows the top responded-to 11 lines followed by lines by indication type. Blank cells represent data not available.

**Supplementary Figure 2.**
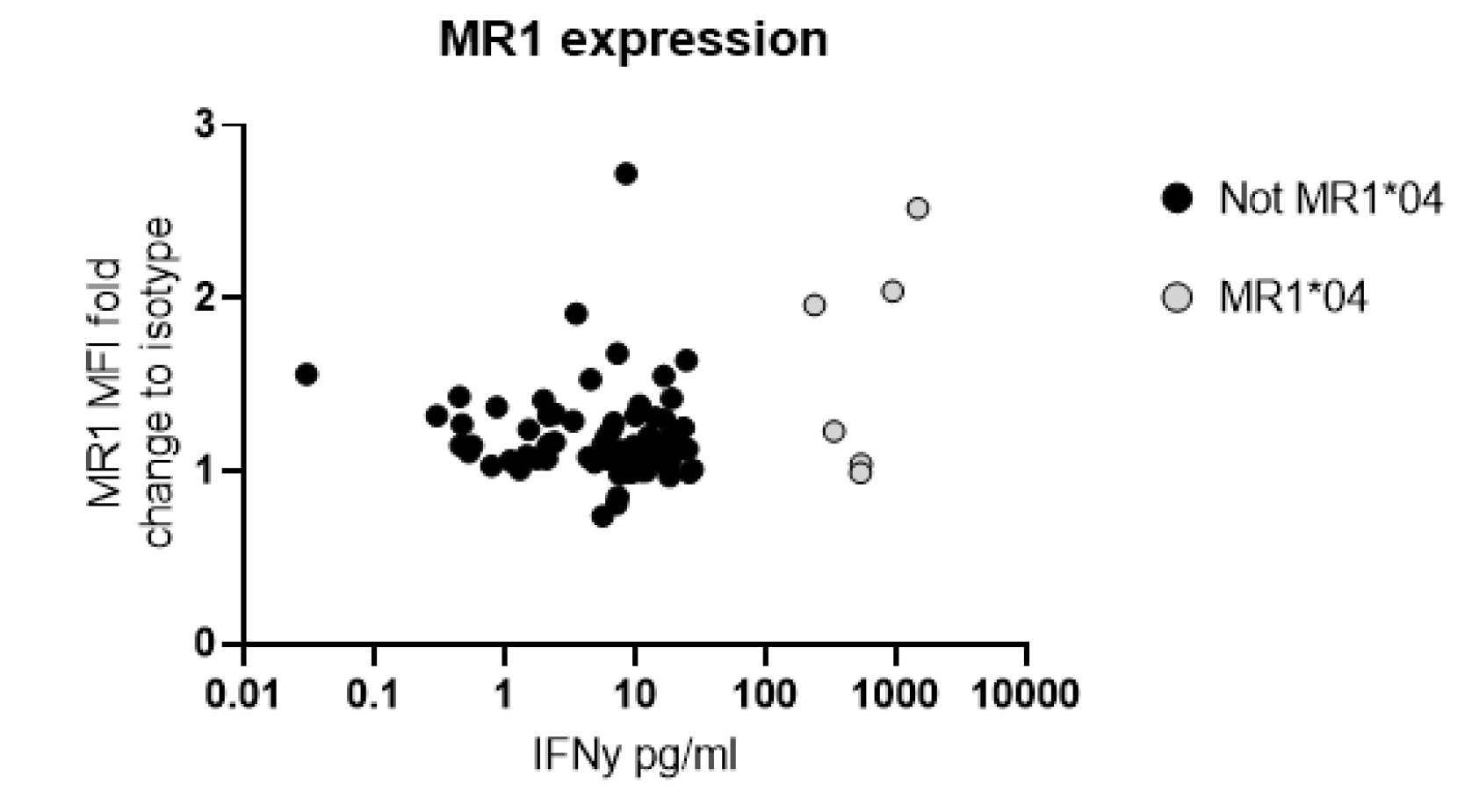
7G5.TCRT killing vs MR1 surface intensity of the target. MR1 expression on cancer cell lines as fold change upregulation compared to isotype control, plotted against TCR-T reactivity data from one donor. Data shown as IFNγ production from T cells following co-culture with cancer cell lines. Each point is an individual cancer cell line. Grey circles are MR1*04 cell lines, and black circles are non-MR1*04.

**Supplementary Figure 3.**
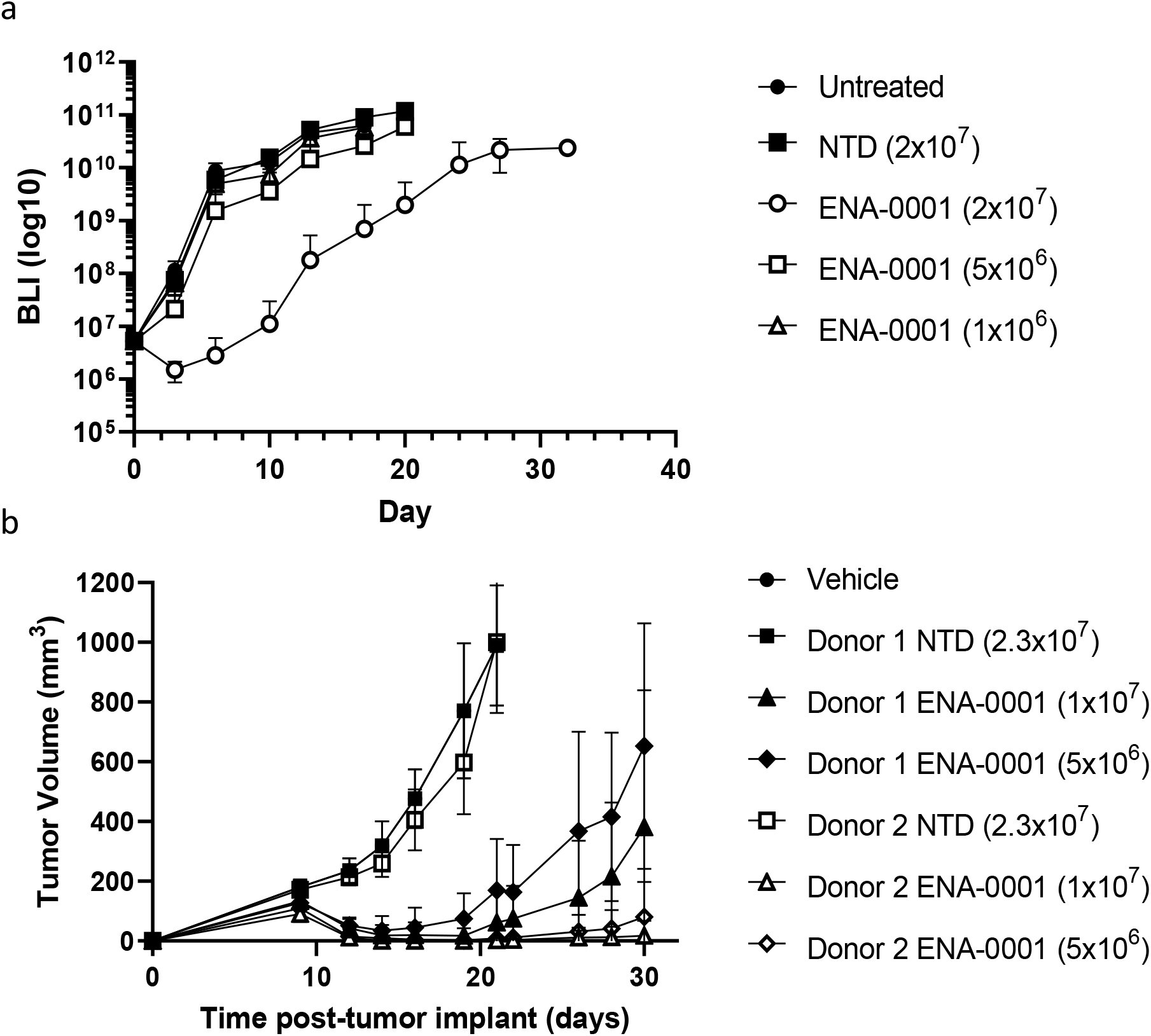
7G5.TCRT effectiveness in murine tumor models **a)** The mice been implanted i.v. with NALM6-Luc tumor cells on day -3 and were then untreated (filled circles), or received 2×10^7^ non-transduced (NTD)T-cells (filled squares) or different doses ranging from 2×10^7^-1×10^6^ of 7G5 TCR T-cells (open symbols as indicated) from one healthy donor on day 0. Mice were injected i.p with luciferin imaged on day 0, day 3 and twice per week thereafter. In this study the T-cells were transduced with a vector that expressed the 7G5 TCR and removed the endogenous TCR using CRISPR-Cas9. Data are shown as the mean +/-SD of whole body Bioluminescence Imaging (BLI) measurements (photons/second) taken from NSG mice (n=5/group). **b)** NSG mice were injected subcutaneously (s.c.) with 5×10^6^ A375-MR1*01 cells. The following day, mice were randomised into groups (n=8/group) and on the same day, were injected i.v. with vehicle (filled circles), or T-cells from donor 1 or 2 (filled and open symbols). These were 2.3×10^7^ total NTD T-cells (squares), or 1×10^7^ (triangles) or 5×10^6^ (diamonds) 7G5 TCR-transduced T-cells. In this study the T-cells were transduced with a vector that only expressed the 7G5 TCR and did not remove the endogenous TCR. Calliper tumour volume measurements were performed three times a week from Day 5. Data are shown as the mean tumour volume (mm3) +/- SD.

**Supplementary Figure 4.**
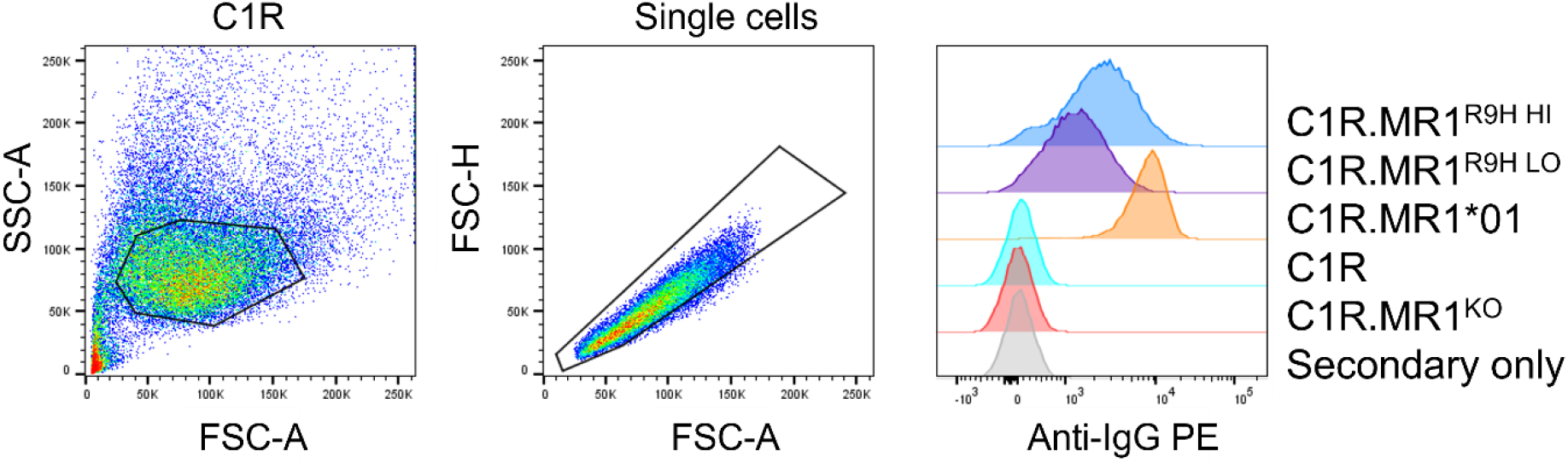
Gating strategy for MR1 expression in C1R derivatives. Gating strategy for MR1 expression of C1R derivatives. Cells were stained with 8F2.F9 hybridoma supernatant, followed by anti-mouse IgG-PE.

**Supplementary Figure 5.**
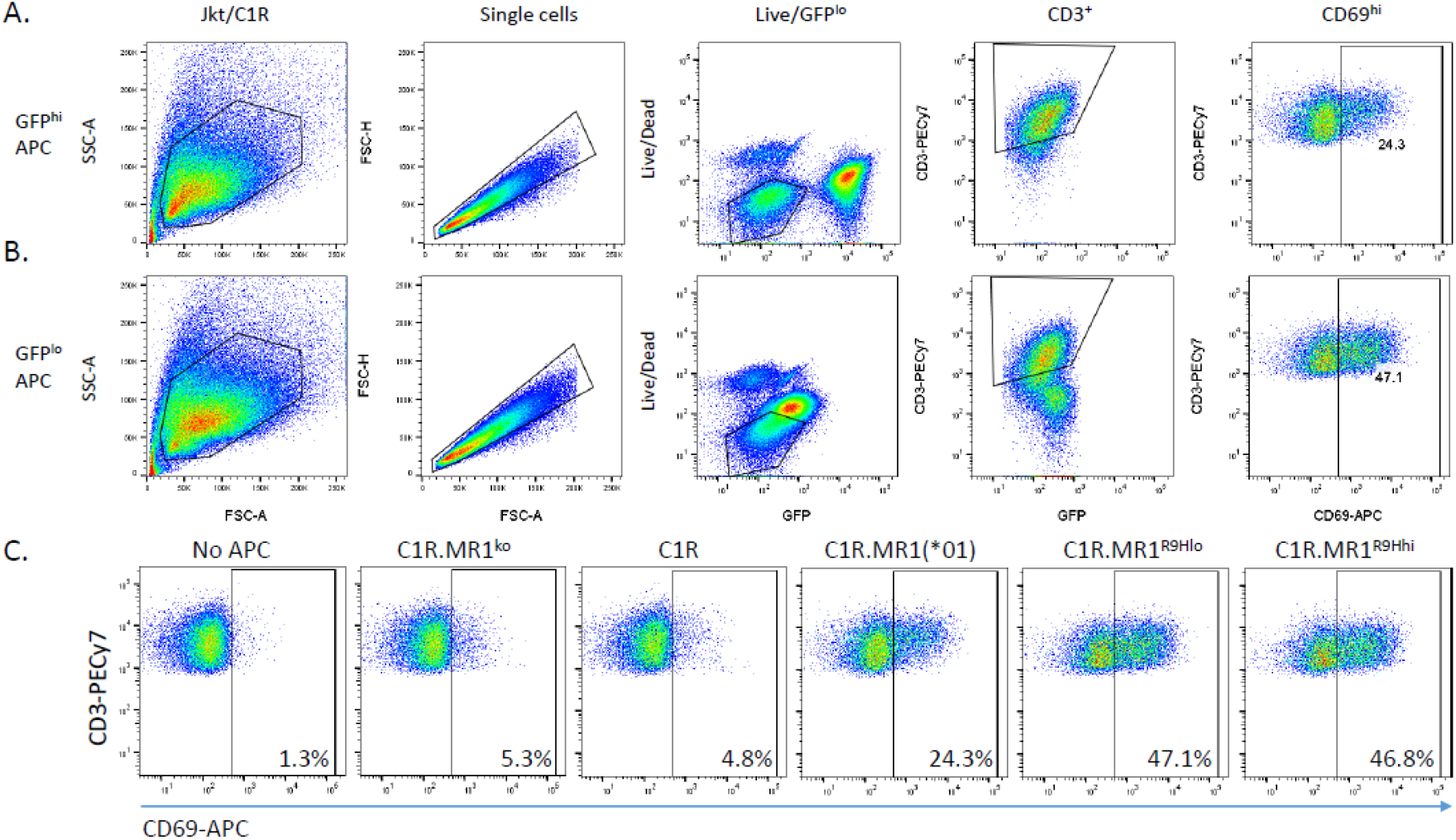
Gating strategy for Jkt.β2m^ko^.7G5 activation assays. Gated for single, live, GFP^lo^, CD3^+^ Jkt.β2m^ko^.7G5 cells prior to assessment of CD69 expression. A) Shows example stimulation with C1R.MR1(*01) cells (GFP^hi^ APC). B) Shows example stimulation with C1R.MR1^R9Hlo^ cells (GFP^lo^ APC). C) Example scatter plots of Jkt.β2m^ko^.7G5 CD69 expression on stimulation with different APCs (left to right: No APC, C1R.MR1^ko^, C1R, C1R.MR1(*01), C1R.MR1^R9Hlo^, C1R.MR1^R9Hhi^).

**Supplementary Figure 6.**
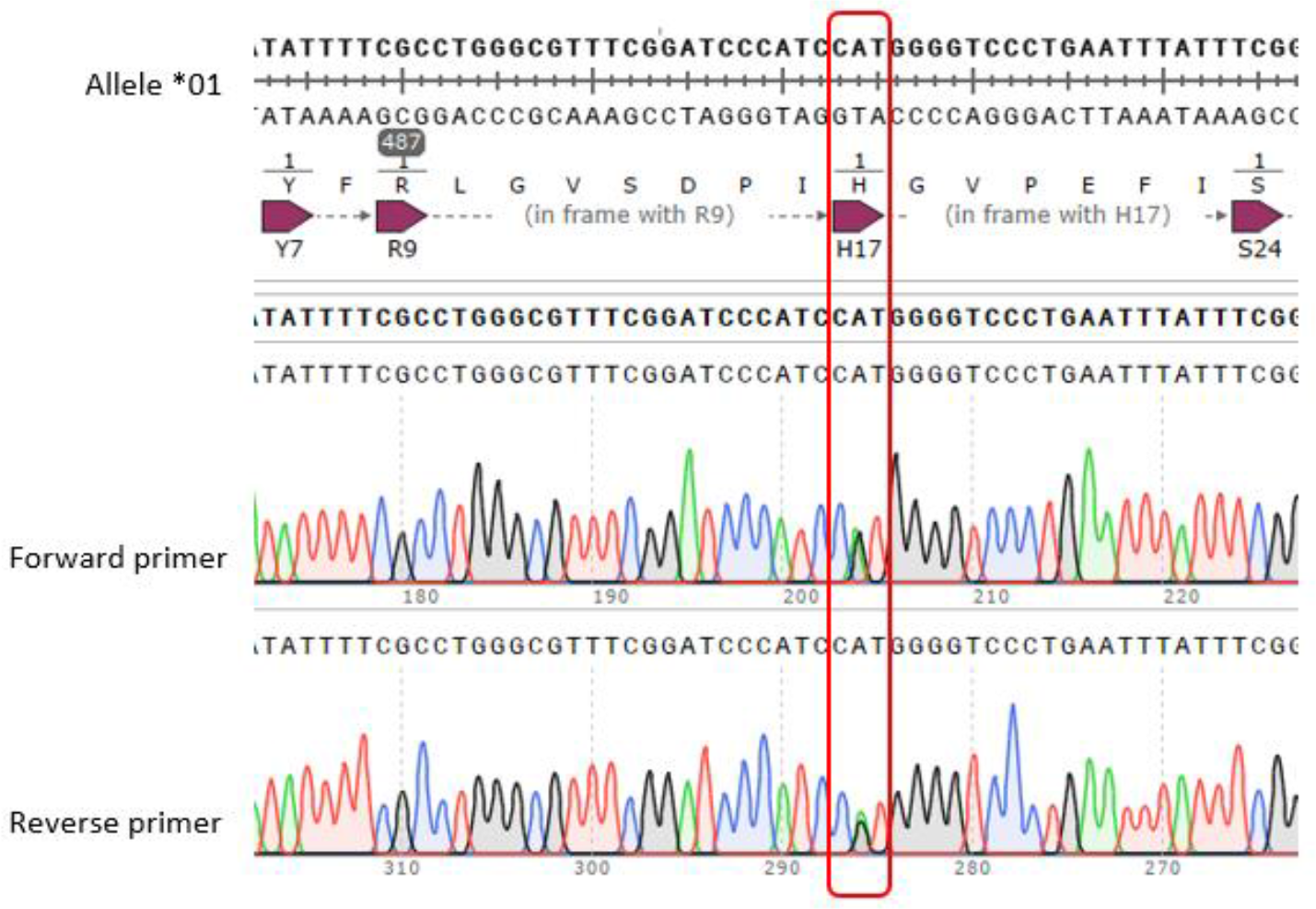
MC.7.G5 clone MR1 sequencing data. Sanger sequencing of the MR1 locus of MC.7.G5 clone analyzed. The MR1*01 allele and MR1*02 allele differ by 1 nucleotide substitution leading to H17R substitution in the MR1*02 allele. The sequences shows the MC.7.G5 clone heterozygous for MR1*01/*02.

